# Ribosomal modification by BUD23 drives selective translational control over energy state and metabolic health

**DOI:** 10.1101/2025.05.16.654455

**Authors:** Noelia Martinez-Sanchez, Anneke Brümmer, Nichola J. Barron, Maria Voronkov, Daniel B. Rosoff, Angélica Liechti, Edward A. Hayter, Sébastien Chamois, René Dreos, Nicolas Guex, Elspeth Johnson, Matthew Baxter, Kerri L.M. Smith, Rebecca C Northeast, Gina Galli, Leanne Hodson, David Gatfield, David W. Ray, David A. Bechtold

## Abstract

Efficient energy metabolism is essential for health and its dysregulation drives cardiometabolic disease. Delivery of regulatory control through translation and ribosome function is emerging as important. Here, we identify the rRNA methyltransferase BUD23 as a potent regulator of cellular and systemic energy homeostasis. Adipocyte-specific deletion of BUD23 in mice regulates lipid and mitochondrial metabolism resulting in a pronounced lean phenotype and resistance to diet-induced obesity. Mechanistically, BUD23 modulates translation initiation and efficiency of mRNAs with specific features – including short 5’ UTR length and GC-rich post-initiation codon usage – characteristic of mitochondrial and lipogenic proteins. Genetic analyses and Mendelian randomisation support a role for BUD23 in human cardiometabolic traits and disease burden. Together, our findings uncover a conserved translational control mechanism that regulates energy state, from cellular metabolism through to human cardiometabolic health.

## Introduction

Despite advances in weight-loss therapies, maintaining long-term energy balance and metabolic health remains a major challenge. Fundamental questions also persist around the molecular and cellular events which set the metabolic programmes of diverse cell types across the body and how they adapt to nutrient excess or deprivation. While transcriptional regulation has been a major focus, growing evidence points to selective control over messenger RNA (mRNA) translation as a key layer of metabolic regulation ^1^.

Ribosomes, once considered uniform protein synthesis machines, are now recognised as compositionally and functionally diverse, enabling the selective translation of mRNA subsets according to cell type and physiological state ^2–5^. Ribosomal RNA (rRNA) is central to this process: it positions mRNA and transfer RNA (tRNA) for accurate codon recognition, ensuring translation fidelity. The small (40S) ribosomal subunit houses the decoding centre, which closely monitors the base pairing between the mRNA codon and the anticodon of the aminoacyl-tRNA to guide correct amino acid incorporation into the growing peptide chain. Eukaryotic rRNA harbours over 200 post-transcriptional modifications ^6^ – predominantly methylations – many at highly conserved and functionally important sites. Some of these modifications are now thought to be dynamic ^7^, raising the possibility that they contribute to ribosomal heterogeneity. However, the functional impact of many such modifications and their potential role in selective translational control remain unclear.

One such modification is N^7^-methylguanosine (m^7^G) at position 1639 of human 18S rRNA (G1575 in yeast), installed by the highly conserved methyltransferase BUD23 (also known as WBSCR22/MERM1) in complex with its obligate dimerization partner, TRMT112 ^8–10^. BUD23 is essential for proper 40S subunit biogenesis and maturation. Although the modification itself has been known for decades, its functional significance for translation has not been determined. In mice, our prior work demonstrated that global *Bud23* deletion causes embryonic lethality, while cardiomyocyte-specific loss results in early postnatal death from cardiac failure, associated with altered mitochondrial function and insufficiency in cardiomyocyte energy production ^11^. These severe phenotypes have until now hindered mechanistic analysis.

Here, we investigate the role of BUD23 in energy metabolism using tissue-specific deletion in adipocytes and hepatocytes – key metabolic cell types. We show that BUD23 is a critical regulator of lipid and mitochondrial metabolism, with distinct effects depending on cellular context. BUD23-sensitive transcripts share characteristic 5′ mRNA features, including short 5′ untranslated regions (UTRs), which limit the presence of upstream open reading frames (uORFs), and high GC content near the translation start codon – properties enriched among nuclear-encoded mitochondrial genes and lipogenic regulators. Structural modelling of 48S pre-initiation complexes suggests that the BUD23-catalyzed m^7^G1639 modification facilitates translation initiation through contacts with initiator tRNA (tRNAi). In line with conservation of function, human genetic analyses and Mendelian randomization link BUD23 expression to cardiometabolic traits, including hepatic lipid content, body mass index, and obesity. Together, our findings identify BUD23 as a key determinant of selective translation with broad relevance to metabolic health and disease.

## Results

### *Bud23* is essential for normal lipid distribution and storage in white adipose tissue

To interrogate the role of BUD23 in regulating energy metabolism in mammals, we generated mice with selective deletion of *Bud23* in white and brown adipocytes using established transgenic lines (*Bud23^fl/fl^* ^11^*; Adipoq^Cre^* ^12^; **Figure S1A,B**).

Under *ad libitum* feeding conditions with standard chow diet, growth and body weight were similar between male mice lacking *Bud23* in adipocytes (*Bud23^fl/fl^;Adipoq^Cre^*, designated herein as Ad^KO^) and their littermate controls (*Bud23^fl/fl^*, designated Ad^WT^; **Figure 1A**). However, analyses of body composition revealed a striking phenotype, wherein Ad^KO^ mice exhibited a pronounced attenuation of fat mass accumulation and a significant increase in lean mass relative to controls (**Figure 1B**). This lean phenotype in Ad^KO^ mice was not associated with altered food intake (**Figure 1C**), nutrient absorption (as judged by faecal energy content; **Figure 1D**), or locomotor activity (**Figure 1E**). We did observe a small, and statistically significant, increase in daytime body temperature in the Ad^KO^ mice (**Figure S1C**). However, housing the animals at thermoneutral conditions (28°C for 6 weeks) failed to normalise genotype differences in fat mass (**Figure S1D-F**), indicating that altered thermoregulation did not underlie the altered adiposity of the Ad^KO^ mice. A similar lean phenotype was observed in female Ad^KO^ animals (**Figure S1G,H**).

**Figure 1.**
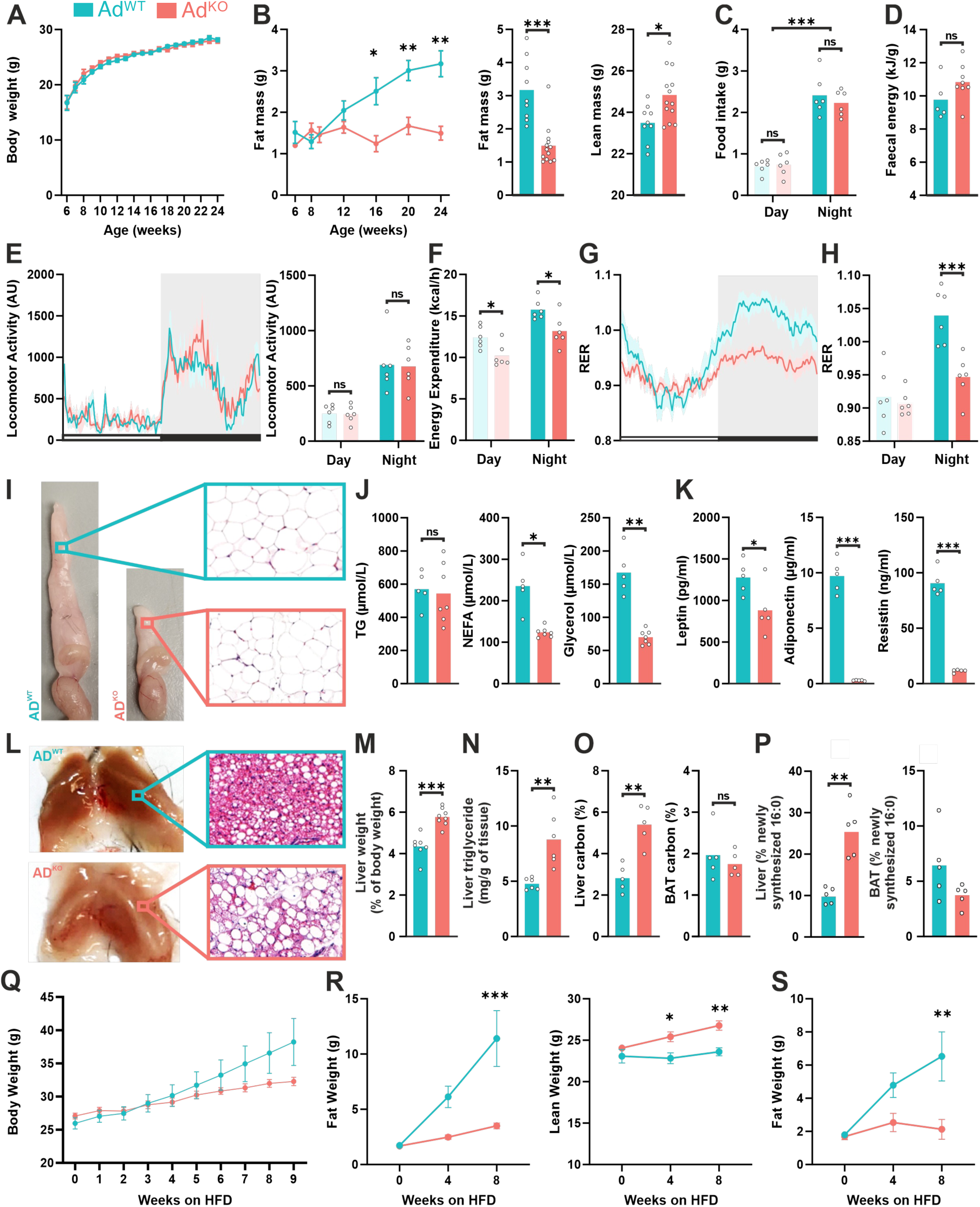
Loss of Bud23 function drives profound metabolic phenotype and shift in body composition. (**A**) Body weight of *Bud23^fl/fl^; Adipoq^Cre+^* (Ad^KO^, n=9) mice compared to littermate controls, *Bud23^fl/fl^* (Ad^WT^, n=11). (**B**) Body composition of Ad^KO^ and Ad^WT^, with total fat mass (left and middle panels) and lean mass (right panel). Histograms reflect body composition at 24 wks. (**C**) Food intake and (**D**) faecal energy content of Ad^WT^ and Ad^KO^ mice. (**E-H**) Locomotor activity (**E**), energy expenditure (**F**) and respiratory exchange ratio (RER; **G,H**) in Ad^WT^ and Ad^KO^ mice (n=6/group). (**I**) Representative images and histology of the gonadal white adipose tissue (gWAT) of Ad^WT^ and Ad^KO^ mice. (**J**, **K**) Circulating serum lipids (triglyceride, NEFA, glycerol) and adipokines (leptin, adiponectin, resistin) in Ad^WT^ and Ad^KO^ mice. (**L**) Representative images and histology of brown adipose tissue (BAT) of Ad^WT^ and Ad^KO^ mice. (**M**, **N**) Liver weight and triglyceride content of AD^WT^ and AD^KO^ mice. (**O**, **P**) Percentage of *de novo* lipogenesis (reflected by incorporation of ^13^C or ^2^H_2_O into palmitate, 16:0) in liver and BAT of Ad^WT^ and Ad^KO^ mice. (**Q**) Body weight of Ad^KO^ and Ad^WT^ in response to 9-week high fat diet (HFD) feeding (n=5/group). (**R**) Fat and lean mass of male Ad^KO^ and Ad^WT^ on HFD. (**S**) Fat mass of female Ad^KO^ and Ad^WT^ mice HFD. *n* reflects biological replicates (mice) in all panels. Mean values are plotted. Statistical significance was tested by two-tailed Student’s t test (B, D, J, K, M, N, O, P) or two-way ANOVA with Holm-Šídák post hoc test (B, C, E, F, H, R, S). ∗p<0.05, ∗∗p<0.01, ∗∗∗p<0.001, ‘ns’ = not significant, based on genotype comparison.

We next profiled metabolic gas exchange and energy expenditure. Despite the lean phenotype, Ad^KO^ mice presented lower daily energy expenditure (**Figure 1F**). Furthermore, the mice exhibited a blunted diurnal profile in respiratory exchange ratio (RER) due principally to a significant reduction in RER at night relative to control mice (**Figure 1G,H**). RER broadly reflects fuel utilisation (carbohydrate oxidation: RER ∼1 vs fatty acid oxidation: RER ∼0.7), and RER values >1 can indicate elevated rates of lipogenesis ^13^. Our findings therefore suggest that the mice lacking BUD23 activity in adipocytes exhibit increased reliance on fatty acid oxidation and, possibly, reduced rates of *de novo* lipogenesis (DNL), both of which would contribute to the reduced fat mass.

Attenuated whole body fat mass in the Ad^KO^ mice was reflected in a significant reduction in subcutaneous and visceral white adipose tissue (WAT) depots, starting at early age and greatly accentuated in older mice (**Figure 1B,I**; **Figure S1I,J**). Interestingly, histological analyses of gonadal WAT (gWAT) sections revealed relatively normal adipocyte morphology in the remaining WAT of these animals (**Figure 1I**, insets; **Figure S1K,L**). Serum triglycerides (TG) were unchanged between genotypes, yet Ad^KO^ presented with significantly lower serum levels of non-esterified fatty acids (NEFA) and glycerol (**Figure 1J**). This suggests that reduced fat storage was not due to excessive lipolysis. To characterize adipose function more broadly, we next assessed circulating adipokine levels. In line with the lean phenotype, leptin, adiponectin, and resistin levels were all reduced in the Ad^KO^ mice (**Figure 1K**), although the magnitude of effect implicates a more direct impact to adipokine production. Together, these findings highlight the profound disturbance in WAT function of Ad^KO^ animals.

Given the significant attenuation of lipid storage in WAT, we next investigated accumulation of lipids within brown adipose tissue (BAT) and liver. Evident at the point of tissue dissection and further confirmed by histological examination, BAT showed accentuated lipid accumulation (“*whitening*”; **Figure 1L**) in the Ad^KO^ mice compared to controls. Similarly, the livers of these mice showed increased weight and TG content (**Figure 1M,N**). These findings suggest that attenuated WAT lipid storage capacity upon loss of BUD23 function drives excess secondary storage in organs such as liver and BAT. Stable isotope labelling with ²H₂O (in drinking water, 48 hr) and [¹³C]-labelled D-glucose (bolus administration following short-term fasting) was used to assess rates of DNL across these tissues. Higher rates of hepatic DNL were confirmed by increased incorporation of both labels into palmitic acid (16:0) in the Ad^KO^ mice relative to controls (**Figure 1O,P**). By contrast, no differences were observed in BAT (**Figure 1O,P**), although local alteration in rates of DNL may be masked by uptake of lipids from the liver. The observed increase in size, TG content, and DNL in the liver underscores a shift in lipid metabolism and storage dynamics in response to adipocyte-specific *Bud23* deletion.

In the Ad^KO^ model, the deletion of *Bud23* occurs in both white and brown adipocytes. However, the phenotypic outcomes described above revealed striking differences, with attenuated and accentuated lipid accumulation in WAT and BAT, respectively. We hypothesised that the whitening of BAT in the Ad^KO^ mice may be secondary to defective lipid storage in WAT. To address this possibility, we generated a *Bud23^fl/fl^*;*Ucp1^Cre^* mouse line (BAT ^KO^), in which *Bud23* is selectively deleted only from brown adipocytes. No differences in body weight, adiposity, or WAT depot weight were apparent in these mice compared with WT littermate controls (BAT^WT^; **Figure S2A-D**). Intrascapular BAT tissue was significantly smaller, but showed no evidence of whitening in the BAT^KO^ mice relative to BAT^WT^ (**Figure S2E**). The animals also showed no overt differences in metabolic or thermogenic phenotypes (**Figure S2F-G**). These analyses highlight that while loss of *Bud23* clearly impacts BAT, the accentuated lipid accumulation in this tissue observed in the Ad^KO^ mice is secondary to WAT dysfunction.

In an attempt to drive lipid accumulation in adipose tissues, we placed Ad^WT^ and Ad^KO^ mice onto high-fat diet (HFD; 60% energy from fat). As expected, control mice gained significant body weight and fat mass over the 9-week HFD treatment (**Figure 1Q-S**). In contrast, Ad^KO^ mice were highly resistant to diet-induced obesity, accumulating significantly less fat mass. These findings demonstrate that adipocyte-specific *Bud23* deficiency impairs WAT lipid storage irrespective of whether the dietary energy source is carbohydrate-rich or fat-rich.

In summary, our results demonstrate that BUD23 is essential for lipid accumulation in WAT. Because adipose tissue is still present in Ad^KO^, the phenotype is unlikely to result from major defects in adipocyte differentiation or survival. Instead, BUD23 appears to play a role in maintaining long-term lipid storage in WAT and preserving mature white adipocyte function.

### *Bud23* is required for white adipocyte function, but not differentiation

We next isolated stromal vascular fraction (SVF) cells from gWAT of Ad^KO^ and Ad^WT^ mice to directly examine adipocyte differentiation capacity *in vitro*. No defects in adipocyte differentiation efficiency, lipid accumulation, or lipid droplet formation in cells derived from Ad^KO^ animals were observed (**Figure S3A,B**). Furthermore, within these newly differentiated adipocytes, expression levels of endocrine hormones and lipid metabolism enzymes were normal (**Figure S3C**). These findings further corroborate that the lean phenotype observed in Ad^KO^ mice *in vivo* is not due to an impaired capacity of adipocytes to differentiate.

Transcriptomics analyses on gWAT isolated from ∼13-week old Ad^KO^ and Ad^WT^ mice (thus prior to the profound differences in fat mass that increase with age) revealed widespread differential gene expression associated with *Bud23* deletion (**Figure 2A**). We first assessed whether cellular composition of WAT contributed to the transcriptional signatures observed. We performed virtual cytometry analyses using CIBERSORT deconvolution ^14^, and leveraging published single-cell RNA-seq data from mouse gWAT ^15^ (**Figure 2B**). While these analyses did not reveal profound changes to cellular composition, including prevalence of adipocyte precursors, notable exceptions emerged. This included a significant reduction in *lipid scavenging adipocyte* (LSA) and *lipogenic adipocyte* (LGA) subpopulations, and a strong increase in *stressed lipid scavenging adipocytes* (SLSA) in Ad^KO^ gWAT (**Figure 2B,C**). These inferred changes in cellular composition align well with the *in vivo* phenotype. LGAs, which are characterised by high expression of genes involved in lipid biosynthesis and insulin responsiveness, likely contribute to DNL and efficient lipid storage ^15^. Their depletion in Ad^KO^ mice, coupled with the concomitant increase in SLSAs, suggests a failure to properly manage and store lipids in the absence of BUD23.

**Figure 2.**
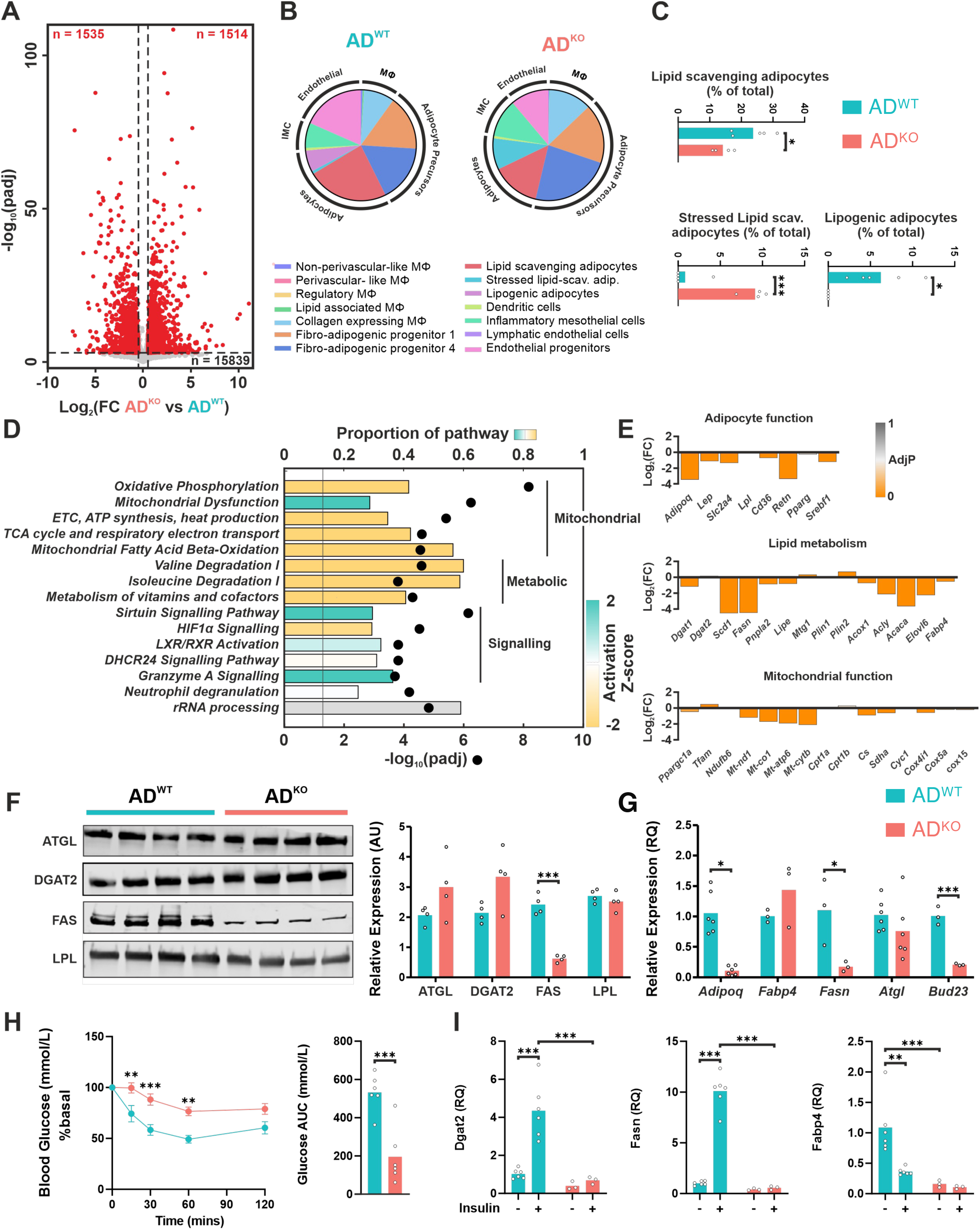
Bud23 is required for normal white adipocyte function. (**A**) Volcano plot of differentially expressed genes from RNA sequencing in Ad^KO^ gonadal white adipose tissue (gWAT) relative to control (n=4-5 mice/genotype). Significantly up-(right) and down-(left) regulated transcripts are indicated in red. (**B**) Virtual cytometry plot displaying inferred cell composition from gWAT RNA sequencing data in Ad^WT^ and Ad^KO^ mice. (**C**) Percentage of *lipid-scavenging adipocytes*, *stressed lipid-scavenging adipocytes* and *lipogenic adipocytes* inferred from gWAT RNA sequencing. (**D**) Pathway enrichment analyses (IPA) of differentially regulated genes in gWAT (black dots reflect *p*-value; columns reflect proportion of pathway, column colour shows predicted pathway activation in Ad^KO^). (**E**) Expression of individual genes related to adipocyte function, lipid metabolism and mitochondrial function (Log2 fold change Ad^KO^ vs Ad^WT^). (**F**) Representative Western blot images (left) and quantification (right) of ATGL, DGAT2, FAS, LPL protein expression in gWAT of Ad^WT^ or Ad^KO^ mice. (**G**) Gene expression in mature adipocytes isolated from gWAT of Ad^WT^ control or Ad^KO^ mice. (**H**) Intraperitoneal insulin tolerance test and area within the curve (AUC) of Ad^WT^ mice and Ad^KO^. (**I**) Insulin responsive gene expression in gWAT explants derived from Ad^WT^ control or Ad^KO^ mice (-= vehicle treatment, + = insulin). n reflects biological replicates (mice) in all panels. Mean values are plotted, ± SEM where appropriate. Statistical significance was tested by two-tailed Student’s t test (C, F, G, H) or two-way ANOVA with Holm-Šídák post hoc test (I). *p<0.05, **p<0.01, ***p<0.001 based on genotype comparison. Panel A, E: adjusted p-value (padj) from RNA sequencing differential expression analysis.

We next undertook pathway enrichment analysis on the gWAT transcriptomics. Ingenuity Pathway Analysis (IPA) revealed a striking downregulation of mitochondrial pathways, broadly affecting key metabolic processes, including respiratory electron transport, ATP synthesis, fatty acid β-oxidation, and branched-chain amino acid (BCAA) metabolism (**Figure 2D**; **Figure S3D**). Increased mitochondrial dysfunction was also predicted. Examination of key differentially expressed genes involved in adipocyte function, lipid metabolism, and mitochondrial activity revealed their extensive downregulation in Ad^KO^ (**Figure 2E**). The reduced expression of adipokine genes, such as *Leptin (Lep)*, *Adiponectin (Adipoq)* and *Resistin (Retn)*, was also consistent with the observed endocrine profiles of Ad^KO^ mice (**Figure 1K**). Importantly, *Bud23* deletion did not cause uniform repression of all adipocyte-specific or metabolic genes. Instead, selective impact to key metabolic pathways, such as fatty acid synthesis, was evident. For example, fatty acid synthase (FAS), a key enzyme in DNL, exhibited a striking decrease in protein abundance in Ad^KO^ WAT, while other markers remained relatively unchanged (**Figure 2F**). Importantly, analyses of mature adipocytes isolated directly from gWAT of Ad^KO^ and Ad^WT^ mice demonstrated that the transcriptomic changes seen in whole tissue RNA-seq originate from *Bud23*-deletion dependent reprogramming of white adipocytes themselves (**Figure 2G**).

Given the well-established influence of insulin over adipocyte function, we assessed insulin sensitivity in Ad^KO^ and Ad^WT^ mice. *In vivo*, clearance of circulating glucose in response to acute insulin administration was significantly attenuated in the knockout mice when compared to controls (**Figure 2H**). Moreover, gWAT explants derived from Ad^KO^ mice cultured *in vitro*, showed a profoundly attenuated induction of *Dgat2*, *Fasn*, and *Fabp4* expression following insulin treatment, compared to the response in Ad^WT^ explants (**Figure 2I**).

We concluded from the above experiments that *Bud23* exerts profound influence over white adipocyte function, yet its effects are selective, with mitochondrial metabolism and lipid biosynthesis as key targets.

### BUD23 influence over lipid metabolism and mitochondrial function is conserved across white and brown adipose

White and brown adipocytes have distinct physiological roles: white adipocytes store and release TGs and fatty acids (FAs) in response to energy demands, whereas brown adipocytes oxidise FAs primarily to generate heat. Given the strong effects of *Bud23* loss on WAT mitochondrial and metabolic programs, we next asked whether similar regulatory patterns occur in BAT, despite its thermogenic specialisation.

As noted above (**Figure S1C-D**), Ad^KO^ mice exhibited relatively normal body temperature profiles, suggesting preserved BAT thermogenic function. Direct measurements confirmed that interscapular BAT temperature was comparable under normal housing (∼22°C ± 2°C; **Figure 3A,B**), and both groups maintained core body temperature during cold exposure (**Figure 3C,D**). Consistent with this, BAT *Ucp1* expression was unchanged at both RNA and protein levels, although several other thermogenesis-associated transcripts differed significantly between genotypes (**Figure S4B,D**). To investigate whether *Bud23* loss reprograms BAT at the molecular level, we performed RNA-seq on BAT from Ad^KO^ and control mice. This revealed extensive differential gene expression (**Figure 3E**), with pathway analyses (IPA) highlighting mitochondrial protein-encoding transcripts and pronounced downregulation of pathways central to oxidative metabolism, including fatty acid oxidation, the tricarboxylic acid (TCA) cycle, the respiratory chain, and BCAA catabolism (**Figure 3F**).

**Figure 3.**
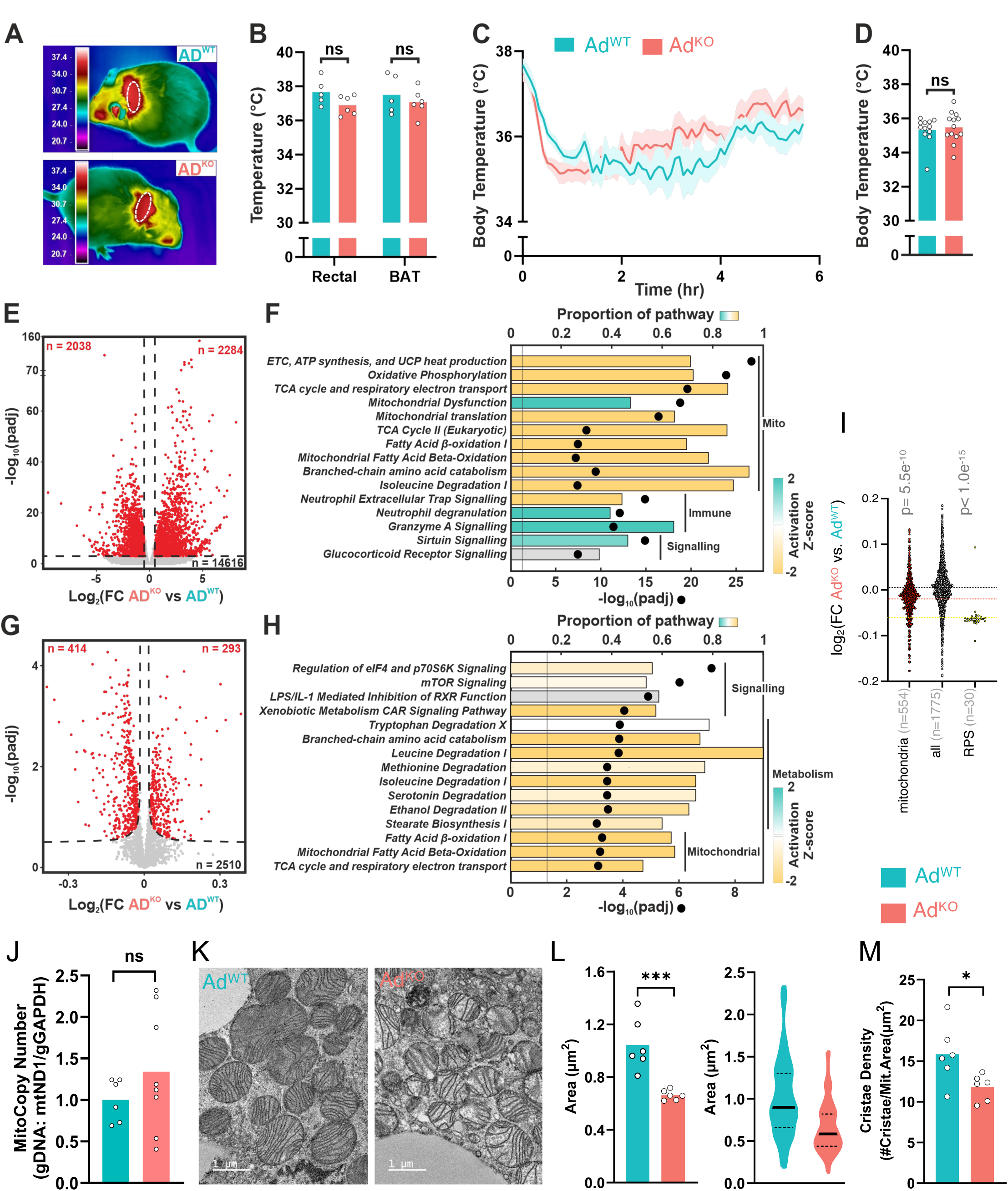
Bud23 directs brown adipose tissue morphology, lipid metabolism and mitochondrial function. (**A**) Representative infrared thermal images of mice (shaved to assess interscapular BAT). (**B**) Rectal and BAT temperature of Ad^WT^ and Ad^KO^ mice (ambient temperature 22 ± 2°C). (**C, D**) Body temperature of Ad^WT^ and Ad^KO^ mice exposed to 4°C for 6 hours. (**E**) Volcano plot of RNA sequencing analyses (n = 8 mice/genotype) highlighting significantly differentially expressed (red) genes in BAT of AD^KO^ relative to Ad^WT^ mice. (**F**) Pathway enrichment analyses (IPA) of differentially regulated genes in BAT (black dots reflect adjusted *p*-value; columns reflect proportion of pathway, column colour shows predicted pathway activation in Ad^KO^). (**G**) Volcano plot of proteomic analyses highlighting differentially expressed proteins in BAT of Ad^KO^ relative to Ad^WT^ mice (n = 8 mice/genotype). (**H**) Pathway enrichment analysis (IPA) of differential proteins expression in response to Bud23 deletion in adipocytes. (**I**) Genotype-specific differences (as log_2_FC, irrespective of padj) in protein abundance of all RPS (small subunit ribosomal factors), mitochondrial associated factors (mito; based on annotated with MitoCarta3.0) and all other proteins detected. (**J**) BAT mitochondria copy number of Ad^WT^ and Ad^KO^ mice. (**K**) Representative electron microscopy images, (**L**) quantification of mitochondrial cross-sectional area (left panel: average/mouse; right panel: all mitochondrion), and (**M**) cristae density (average/mouse). n reflects biological replicates (mice) in all histograms. Mean values are plotted (with ± SEM where appropriate). Statistical significance was tested by unpaired two-tailed Student’s t test (B, D, J, L) or one-way ANOVA with Dunnet’s post hoc test (I). ***p<0.001, ns = not significant, based on genotype comparison. Panel E, G and I: log₂FC (fold change) vs. –log₁₀(adjusted *p*-value) from differential expression analysis.

To determine whether these transcriptomic changes were reflected at the protein level, we carried out proteomic profiling (**Figure 3G**). This confirmed widespread differential protein expression, with IPA revealing enrichment for mitochondrial proteins and BCAA metabolic enzymes, including marked reduction of BCAT2 (**Figure 3H**, **Figure S4D**). Notably, proteomics also identified two translation-associated pathways – *Regulation of eIF4 and p70S6K Signaling* and *mTOR Signaling* pathways – as significantly dysregulated. These pathways share overlapping components, including ribosomal proteins, translation initiation factors, and key signalling components that integrate hormonal, nutrient, and stress cues to regulate both global and transcript-specific translation. Gene ontology analyses reinforced these findings, revealing reduced abundance of proteins associated with mitochondrial function, lipid metabolism, and ribosomal subunit biogenesis (**Figure S4C**). Small ribosomal subunit proteins (RPS) were particularly affected (**Figure 3I**), consistent with BUD23’s established role in 18S rRNA modification and small subunit maturation ^8,16,17^.

Given the consistent transcriptomic and proteomic signatures of mitochondrial disruption, we next examined BAT mitochondrial content and ultrastructure. Mitochondrial DNA copy number was unchanged (**Figure 3J**), but electron microscopy (EM) revealed reduced mitochondrial area (**Figure 3K,L**) and diminished cristae density in *Bud23*-deficient BAT (**Figure 3K,M**). Proteomic data also pointed to depletion of proteins governing mitochondrial dynamics and inter-organelle contacts (e.g. MFN1/2, OPA1, MIGA2), alongside changes in lipid droplet coat proteins (e.g. PLIN2, PLIN5) and mitochondrial import machinery (e.g. TIMM22) (**Figure S4D**). These alterations occurred without compromising thermogenic performance, suggesting that BUD23 loss selectively perturbs mitochondrial organization and its integration with lipid storage and metabolism.

In summary, *Bud23* deletion in adipocytes induces convergent molecular defects in WAT and BAT, characterised by downregulation of mitochondrial, lipogenic, and ribosomal protein networks. The proteomic shifts are consistent with impaired translation due to loss of 18S rRNA methylation, while the broader transcriptional and metabolic reprogramming likely reflects a combination of primary and secondary effects. Together, these findings position BUD23 as a key regulator of mitochondrial architecture and metabolic function across adipose depots.

### Hepatic targeting of *Bud23* reveals selective impact to energy homeostasis and mitochondrial metabolism

Due to its relatively homogenous cellular composition, large size, and high metabolic activity, we considered the liver well-suited for investigating the functional impact of BUD23 on translation. To this end, we generated a hepatocyte-specific *Bud23* deletion by crossing *Bud23^fl/fl^*mice with the well-established inducible *Alb^CreERT^*^2^ line ^18^. Tamoxifen administration to *Bud23^fl/fl^*;*Alb^CreERT^*^2^ (Liv^KO^) and control *Bud23^fl/fl^* (Liv^WT^) animals resulted in a robust reduction in *Bud23* expression in Liv^KO^ animals (**Figure S5A,B**).

We first profiled the Liv^KO^ model for overt metabolic phenotypes. Under standard conditions, no significant differences between Liv^KO^ and Liv^WT^ mice were observed for body weight, body composition, energy expenditure, RER, or food intake (**Figure 4A-F**). Interestingly, despite this overall metabolic similarity, hepatic TG levels were elevated in Liv^KO^ mice (**Figure 4G**). We next imposed, metabolic challenges driving either positive or negative energy balance (HFD-feeding and fasting, respectively) to further examine *Bud23*-dependent metabolic phenotypes. During 8 weeks of HFD feeding, Liv^KO^ and Liv^WT^ mice showed similar body weight gain (**Figure 4H**) and insulin response (**Figure S5D**), but with elevated levels of hepatic TG in Liv^KO^ (**Figure 4I**). When subject to an extended fast (48 h), Liv^KO^ mice exhibited significantly attenuated β-hydroxybutyrate production, despite showing similar levels of body weight loss and hypoglycaemia (**Figure 4J-L**). Both findings are consistent with altered mitochondrial activity (e.g. β-oxidation, BCAA) and lipid handling.

**Figure 4.**
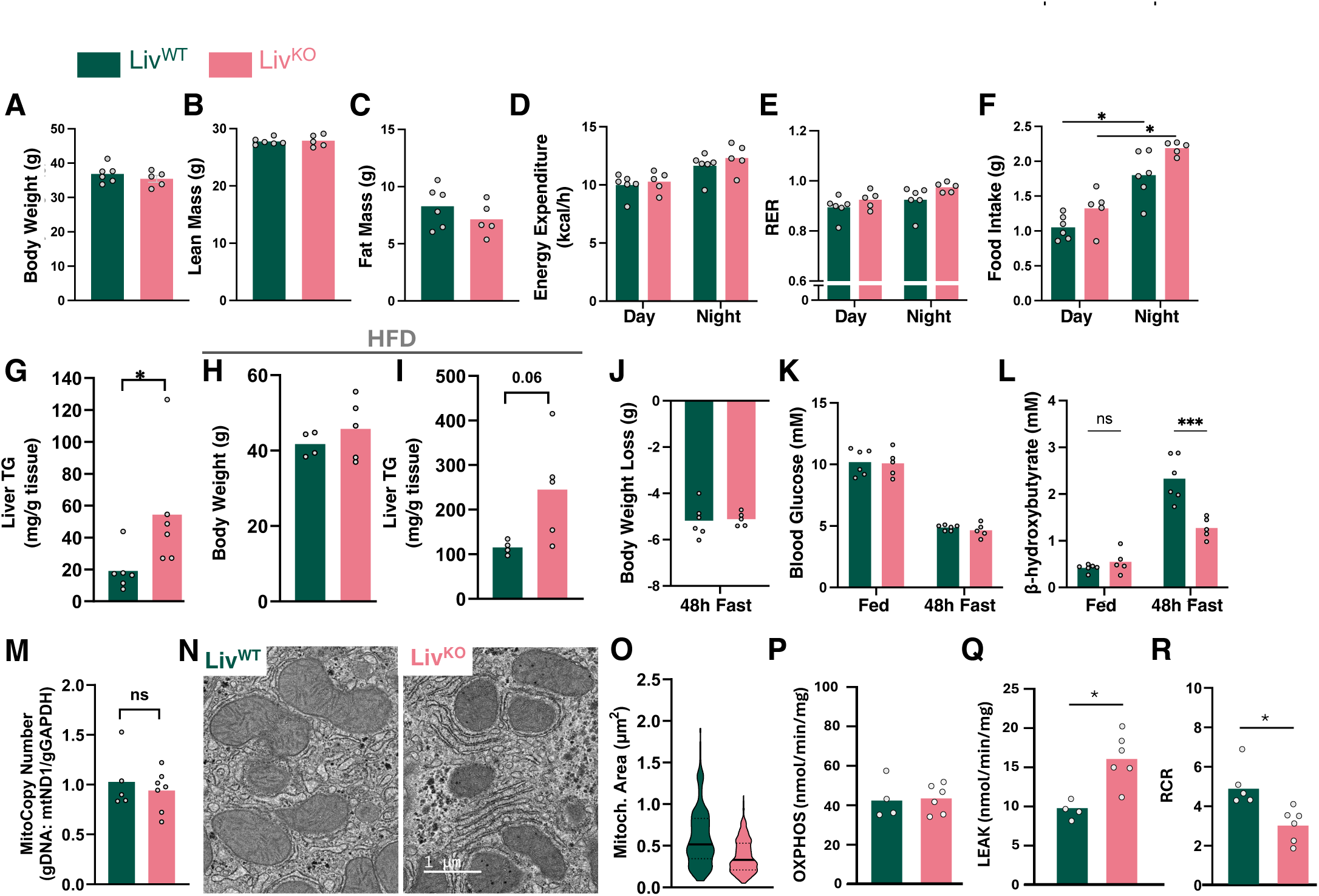
Bud23 directs a selective impact to liver function and proteome. (**A-G**) Body weight (**A**), lean mass (**B**), fat mass (**C**), energy expenditure (**D**), respiratory exchange ratio (RER) (**E**), food intake (**F**), and hepatic triglyceride levels (**G**) of Liv^WT^ and Liv^KO^ mice at 4 weeks post-tamoxifen induced recombination. (**H, I**) Body weight (**H**) and hepatic triglyceride levels (**I**) of Liv^WT^ and Liv^KO^ mice in response to 8 weeks of high fat diet (HFD) feeding. (**K-L**) Body weight loss (**K**), blood glucose (**J**) and circulating β-hydroxybutyrate (**L**) in Liv^WT^ and Liv^KO^ mice in response to a 48-hour fast. (**M**) Mitochondria copy number in liver of Liv^WT^ and Liv^KO^ mice. (**N**) Representative electron microscopy images and (**O**) quantification of mitochondrial cross-sectional area. (**P-R**) OROBOROS mitochondrial respiration rate analysis of isolated mitochondria from liver of Liv^WT^ and Liv^KO^ (n=4-6). OXPHOS = state III respiration; LEAK = state IV respiration; RCR = respiratory control ratio. *n*, reflects biological replicates. Mean values are plotted. Statistical significance was tested by unpaired two-tailed Student’s t test (A-C, G, H, I, K, M, O-R) or two-way ANOVA with Holm-Šídák post hoc test (D-F, J, L). ∗p<0.05, ∗∗∗p<0.001, ‘ns’ not statistically significant.

Given our findings in WAT and BAT, we examined the impact of *Bud23* loss on mitochondrial content, structure and function in the livers of Liv^KO^ and Liv^WT^ mice. Whole tissue mitochondrial copy number was unchanged (**Figure 4M**) and EM analyses revealed a small reduction in mitochondrial area in Liv^KO^ (**Figure 4N,O**). We therefore isolated hepatic mitochondria and assessed potential genotype differences in respiration using the OROBOROS platform (**Figure 4P-R**). Respiration rates measured in isolated mitochondria in the presence of complex I substrates and saturating levels of ADP (State III respiration; OXPHOS) were comparable between genotypes (**Figure 4P**). However, a significant elevation in State IV respiration (LEAK) reflecting proton leak-driven respiration was evident in mitochondria isolated from Liv^KO^ mice (**Figure 4Q)**. This in turn resulted in a significantly reduced respiratory control ratio (RCR) (**Figure 4R)**, strongly suggesting that mitochondrial ATP production is less efficient in Liv^KO^ tissue.

Collectively, our findings demonstrate that mitochondrial dysfunction and lipid metabolism alterations are common features of *Bud23* deficiency across tissues.

### Hepatic *Bud23* loss selectively remodels translation

We next took advantage of the liver as a platform to dissect the molecular mechanism of BUD23 action at the level of mRNA translation. To this end, we performed transcriptomic (RNA-seq) and ribosome profiling (Ribo-seq) on livers from Liv^KO^ and Liv^WT^ animals at an early stage after gene deletion (3 weeks post-induction). Given the well-documented diurnal rhythmicity of hepatic metabolic activity, gene expression, and translation ^19^, we collected samples in fasted (*Zeitgeber* Time ZT6; mid-light phase) and fed (ZT18; mid-dark phase) states (**Figure S6A)**. Principal component analyses (PCA; **Figure S6B-D**) revealed that genotype (PC1) was the main driver of variance for both mRNA abundances and ribosome occupancies, followed by time-of-day effects (PC2). Translation efficiency (TE) – defined as ribosome occupancy signal normalised to RNA abundance i.e. Ribo-seq/RNA-seq – also showed a strong genotype effect, yet little time-of-day contribution. Therefore, unless stated otherwise, subsequent analyses were carried out on combined timepoint data.

*Bud23* loss broadly altered mRNA abundances and ribosome occupancies, with similar numbers of genes up– and down-regulated (**Figure 5A,B**). Changes in TE were more selective (105 decreased; 179 increased; **Figure 5C**). GO analysis highlighted two main trends: (i) increased abundance and TE for ribosomal protein (RP) and translation factor mRNAs, consistent with a compensatory response to reduced small ribosomal subunit levels reported for *Bud23*-deficient cells ^8^; and (ii) decreased mRNA and footprint abundance for metabolic and mitochondrial genes (**Figure 5D**). Several key mitochondrial and lipid metabolic transcripts (e.g. *Agpat2, Agpat3, Ak3, Bola3, Coa5, Rxra, Sirt3*) were also significantly reduced in their TE (**Figure S6E**). However, as the overall number of significant TE-affected genes was relatively small, conventional GO *overrepresentation* analyses did not highlight mitochondrial categories (**Figure 5D**). To better capture coordinated trends, we applied a complementary GO enrichment analysis that tests for systematic TE shifts across gene groups rather than relying on significant genes only. This analysis revealed that metabolic and mitochondrial gene sets were among the most consistently reduced in TE, whereas cytoplasmic translation-related sets (notably ribosomal protein mRNAs) dominated the TE-increased group (**Figure 5E**). Many ribosomal protein– and translation-related transcripts bear 5’ terminal oligopyrimidine (TOP) motifs regulated by mTOR ^20^. In Liv^WT^ controls, TE of TOP mRNAs rose significantly from ZT6 to ZT18, consistent with feeding-dependent mTOR activation, and as observed previously ^19,21,22^ (**Figure 5F**). In contrast, Liv^KO^ samples exhibited constitutively high TE at both timepoints for these transcripts, thereby abolishing the normal rhythmic pattern and indicating that the proper coupling of TOP mRNA translation to feeding state requires BUD23 activity.

**Figure 5.**
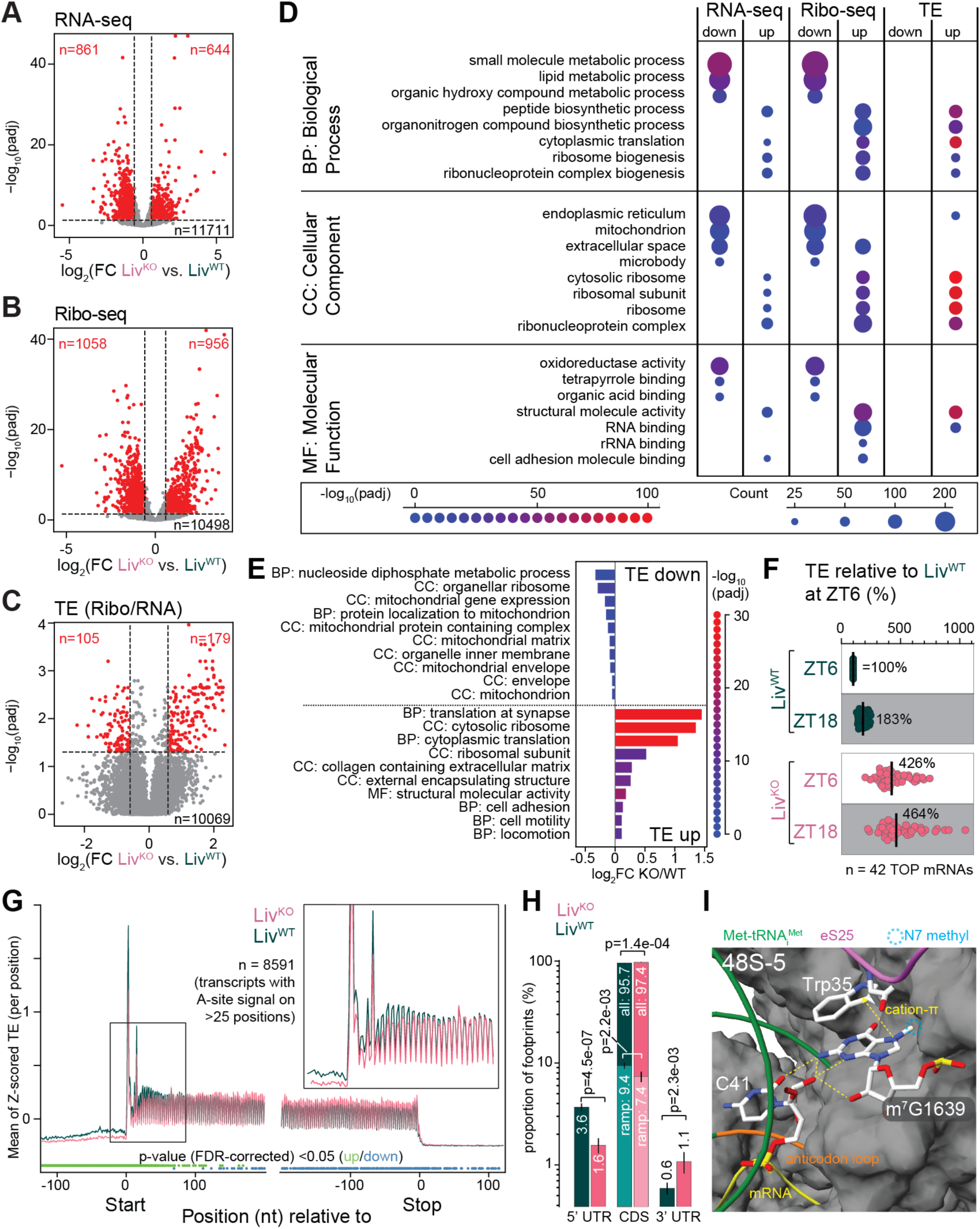
Hepatic *Bud23* drives selective impact on translation efficiency. (**A**) Volcano plot of differentially expressed genes in liver from RNA sequencing in Liv^KO^ vs Liv^WT^. Significantly up-(right) and down-(left) regulated transcripts are indicated in red (adjusted p-value <0.05, log_2_FC >1.5; n = 6 per genotype combined for ZT6 and ZT18 timepoints). (**B**) Volcano plot as in (**A**) but visualising differential Ribo-seq data. (**C**) Volcano plot as in (A) but visualising differential translation efficiencies (TE, ratio of normalised Ribo-seq to RNA-seq reads calculated per liver sample). (**D**) GO-term analysis on differentially expressed gene sets shown in panels A-C. Most significant enrichments are shown. Size of circles indicates the number of genes within a category; colour coding indicates statistical significance, as illustrated in the legend in lower part of panel. (**E**)) GO term analysis comparing the fold changes (KO/WT) for genes within a GO set with the fold changes of all genes (using all GO terms with at least 20 genes). Top 10 most significant down– and up-regulated terms are shown. Terms have been sorted according to values of median log2FC (gene set) – median log2FC for all genes in KO vs. WT. (**F**) Translation efficiency in Liv^KO^ (pink) and Liv^WT^ (dark green) at ZT6 (light phase) and ZT18 (dark phase) of selected ribosomal protein/translation factor mRNAs (n=42) that contain a 5’-TOP sequence and previously identified as rhythmic in their TE {Janich, 2015 #2087}. For each transcript, TE is expressed relative to the level in Liv^WT^ at ZT6, which was set to 100%. (**G**) Metagene plot aligning ribosome footprint A-sites relative to coding sequence (CDS) start (left) and stop (right) codons, for Liv^KO^ (pink) and Liv^WT^ (dark green). Transcripts are included with A-sites at 25 or more positions within the shown range of nucleotides (n=8591). Dots below the plot indicate positions with a ratio between A-sites for Liv^KO^ and Liv^WT^ significantly higher (green) or lower (blue) than 0 calculated using a directional one-sample *t*-test. The inset at the top right shows a zoom on the area around the start codon, indicating a specific decrease of ribosomes in Liv^KO^ over the first ∼20 codons and the 5’ UTR. (**H**) Quantification of global read distribution in Liv^KO^ (pink) and Liv^WT^ (dark green) for 5’ UTR, CDS (all and first 20 codons in light colours) and 3’ UTR. Error bars indicate the standard deviation across the 6 samples per genotype. P-values indicate statistical significance assessed by a two-tailed independent *t*-test. (**I**) Structural model of human late initiation complex 48S-5 (PDB 8pj5) at the site of m^7^G1639. Highlighted residues apart from m^7^G1639 (with N^7^-methyl group marked by blue halo) are Met-tRNA_i_^Met^ (backbone traced in green; anticodon loop in orange), tRNA residue C41 that forms extensive hydrogen bonds with m^7^G1639. Trp35 of eS25/RPS25 is placed for cation-π stacking onto m^7^G1639. mRNA trace is marked in yellow and 18S rRNA surface (excluding m7G1639 to see its interactions) in grey.

For TE calculation, ribosome footprint reads are summed over the entire coding DNA sequence (CDS). However, local variation in ribosome occupancy across the CDS can provide insights into ribosome elongation dynamics and codon specificity. We therefore assessed translation elongation by calculating ribosome dwell times for specific codons at the ribosomal E-, P-, and A-sites, based on transcriptome-wide codon occupancies ^23^. While known differences in decoding speeds across codons were evident, these were only minimally modulated by genotype (**Figure S6F,G**), suggesting that elongation dynamics are largely preserved in the *Bud23* knockout. Next, we used metagene analyses to examine the distribution of ribosome footprints around translation start and stop codons (**Figure 5G**). This analysis revealed two striking translation initiation-related alterations in Liv^KO^: (i) significantly reduced ribosome occupancy over 5′ UTRs (**Figure 5G, H**), reflecting loss of upstream ORF translation ^19^; and (ii) reduction of the characteristic “translational ramp” (**Figure 5G, inset**), a region of elevated ribosome density at the beginning of the CDS that gradually declines as elongation progresses. Reduced 5′ UTR occupancy and loss of the ramp in Liv^KO^ both suggest a disruption in early ribosome dynamics, hinting at BUD23 action on translation initiation rather than elongation.

To understand the mechanism of how BUD23-dependent modification at m^7^G1639 can impact translation initiation, we examined cryo-EM structures capturing key intermediates of the human initiation pathway ^24^. In these structures, m^7^G1639 is located in immediate vicinity to the P-site-bound initiator-tRNA (Met-tRNA_i_^Met^), interacting with cytosine 41 (C41) of its anticodon stem loop via hydrogen bonding (**Figure 5I, Figure S7**). The C41 residue is critical for maintaining the unique conformation of initiator-tRNA, distinguishing it from elongator tRNAs, and ensuring proper start codon recognition ^25^. The N^7^-methyl group of m^7^G1639 fills a hydrophobic pocket that helps stabilize this interaction, while the positive charge conferred onto the guanosine ring by the methyl group (quaternary nitrogen at N7) enables cation-π stacking with Trp35 of ribosomal protein eS25/RPS25, particularly in the late initiation complex 48S-5 (**Figure 5I**). This interaction likely further cradles G1639 in an optimal orientation for initiator-tRNA placement. In the absence of m^7^G1639 methylation (i.e., *Bud23* knockout), it is likely that weakened interaction at this site would alter the precision and stability of initiator-tRNA engagement at the start codon.

In summary, BUD23 loss produces selective translational changes – upregulation for ribosomal protein mRNAs and downregulation on mitochondrial protein transcripts – that are associated with altered translation initiation rather than elongation.

### Transcript architecture, including GC-rich ramp region and short 5′ UTRs, dictates BUD23-dependent translation

To identify sequence features underlying BUD23-dependent translation, we first calculated individual TE values for distinct transcript regions: the 5′ UTR, the CDS, and the first 60 nt of the CDS downstream of the start codon (“ramp region”; normalized to total CDS TE and hereafter referred to as ‘normalised Start TE’) (**Figure 6A**). Fold-changes in TE between Liv^KO^ and Liv^WT^ correlated only weakly across these regions (**Figure S8A-E**), indicating that BUD23 is likely to impact them through independent mechanisms.

**Figure 6.**
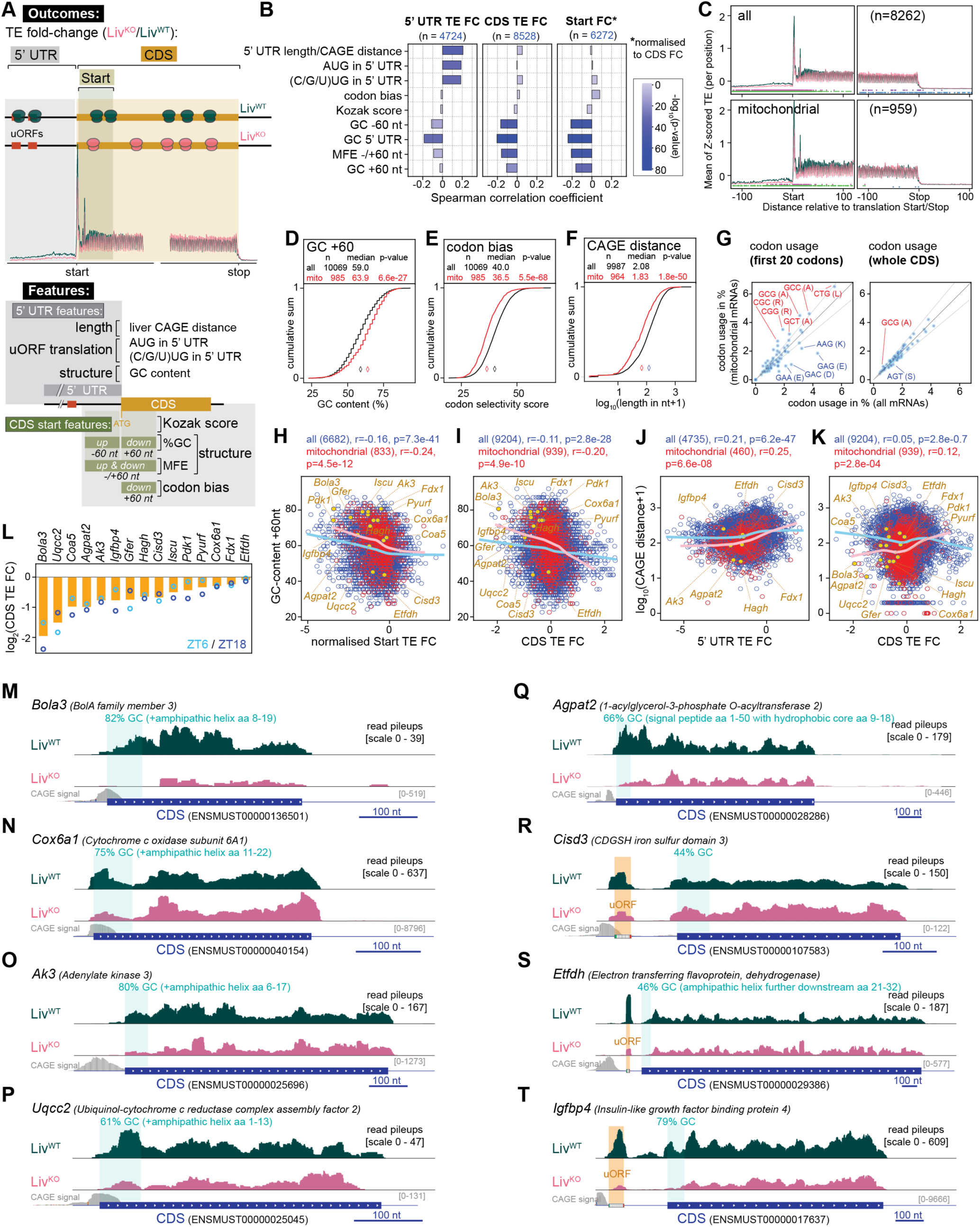
Selective BUD23 regulation is associated with specific transcript features. (**A**) Schematic of analysis of correlation between transcript features and BUD23-selective regulation. Three “Outcomes”, shown at the top of figure, were extracted from the ribo-seq profiles and quantified transcriptome-wide – translation efficiency (TE) change on the 5’ UTR (indicative of uORF translation), TE change on the whole CDS (i.e. change in overall translation rate) and specifically within the first 20 codons (the “ramp effect”). These were correlated with “Features” of transcripts, pertaining to the 5’ UTR – UTR length as determined from cap analysis of gene expression (CAGE) data, presence of AUG-or (C/G/U)UG-initiated uORFs, GC content – or the CDS initiation codon environment – Kozak score, GC content, minimal free energy of folding (MFE), codon bias. (**B**) Spearman correlation coefficients between transcript sequence features (y-axis) and TE log_2_FC (Liv^KO^ vs Liv^WT^) at 5’ UTRs (first panel), CDSs (second panel), and Start codon regions (first 20 codons), as well as the whole CDS (third panel). ZT6 and ZT18 timepoints were combined for FC analyses. (**C**) Metagene plot aligning ribosome footprint A-sites relative to coding sequence (CDS) around start (left panels) and stop (right panels) codons for all transcripts (upper panels) and transcripts encoding mitochondrial proteins (based on MitoCarta 3.0; lower panels) for Liv^KO^ (pink) and Liv^WT^ (dark green) Ribo seq. See Fig. 5G for additional information. (**D**) Analysis of GC content within the 60 nt after the start codon, for transcripts encoding mitochondrial proteins (red) vs all transcripts (black). In the upper part of the panel, the number of transcripts used for the analysis (n), their median (also shown as diamond symbol in graph) and the p-value for the difference using a ranksum test are given. (**E**) As in (D), for codon bias/selectivity for the first 20 codons. Mitochondrial groups composed of significantly less abundant codons. (**F**) As in (D), for 5’ UTR length as determined from cap analysis of gene expression (CAGE) data. (**G**) Analysis of codon composition of first 20 codons (left panel) and whole CDS (right panel), comparing all mRNAs (x-axis) with mitochondrial protein mRNAs (y-axis). Codons are labelled with a 20% higher (red) or lower (blue) abundance in mitochondrial protein mRNAs than in all mRNAs. (**H**) Scatter plot of TE log_2_FC (Liv^KO^ vs Liv^WT^) at the first 20 codons relative to whole CDS, and GC-content in the same +60 nt region of the transcript. All transcripts and mitochondrial mRNAs are shown as blue and red circles, respectively. The local regression fits (LOESS) for all and mitochondrial transcripts are shown in pale blue and pink, respectively. Spearman correlation coefficient and *p*-value of correlation are given above, alongside number of transcripts used in the analysis. Specific genes/transcripts are highlighted. (**I**) As in (H), but showing correlation between whole CDS TE fold-change (Liv^KO^ vs Liv^WT^) and GC-content of +60 nt region. (**J**) As in (H), but showing correlation between 5’ UTR TE FC and 5’ UTR length/CAGE distance. (**K**) As in (H), but showing correlation between CDS fold-change and 5’ UTR length/CAGE distance. (**L**) Quantification of CDS TE change for selected transcripts labelled in panels H-K. Values for ZT6 and ZT18 timepoints are plotted individually as circles, with orange bars showing the mean across both timepoints (n=6). (**M**) Genome browser tracks showing read pile-up along *Bola3 (BolA family member 3)* transcript in Liv^WT^ (upper track, dark green) and Liv^KO^ (lower track, pink). Grey track shows CAGE signal from mouse liver FANTOM5 data (i.e. this 5’ UTR is extremely short). Transcript model and scale bar shown in blue. Green shading placed on first 20 codons, with GC content (82%) and aa position of amphipathic helix written above. Data from ZT18 are plotted, combining reads from the three biological replicates. (**N**) As in (M) for *Cox6a1 (Cytochrome c oxidase subunit 6A1)*, which has very high GC content and very short 5’ UTR. (**O**) As in (M) for *Ak3 (Adenylate kinase 3)*, which has very high GC content and relatively short 5’ UTR. (**P**) As in (M) for *Uqcc2 (Ubiquinol-cytochrome c reductase complex assembly factor 2)*, which has average GC content yet very short 5’ UTR. (**Q**) As in (M) for *Agpat2 (1-acylglycerol-3-phosphate O-acyltransferase 2)*, which has average GC yet very short 5’ UTR (32 nt according to CAGE data). (**R**) As in (M) for *Cisd3 (CDGSH iron sulfur domain 3)*. The position of a uORF is indicated by orange shading. (**S**) As in (R) for *Etfdh (Electron transferring flavoprotein, dehydrogenase)*. (**T**) As in (R) for *Igfbp4 (Insulin-like growth factor binding protein 4)*.

We then quantified a panel of transcript features potentially linked to these TE outcomes (**Figure 6A**, lower). For the 5′ UTR: length (from mouse liver CAGE data); number of translated uORFs (including AUG and near-cognate start codons); and GC content. For the start codon environment: Kozak score; GC content in the –60 nt and +60 nt windows around the AUG; RNA folding minimum free energy (MFE) for the ±60 nt region; and codon bias in the first 20 codons (lower scores = more rare codons). Several associations emerged (**Figure 6B, Figure S8F,G,P**): (i) 5′ UTR TE fold-change correlated positively with UTR length and uORF number, but negatively with UTR GC content; (ii) Reduced CDS TE and normalised Start TE in Liv^KO^ were strongly associated with high GC content and with stable predicted secondary structure (low MFE) near the start codon; (iii) Kozak score and codon bias showed little correlation with TE changes. Because many of these features are intrinsically correlated (e.g. GC content in the UTR and ramp; UTR length and uORF count; **Figure S8H**), we applied a multiple linear regression model. This identified 5′ UTR GC content as the strongest independent predictor for TE change (**Figure S8I**), with additional contributions from UTR length, MFE, and others.

### Mitochondrial transcripts are enriched for BUD23-sensitising features

We next asked whether these features explain the downregulation of many mitochondrial transcripts in *Bud23* knockout tissues (**Figure 5E**, **Figure 6C**). Indeed, nuclear-encoded mitochondrial mRNAs were strongly biased towards BUD23-sensitising features (**Figures 6D-F, S8J-O**). In particular, GC content in the +60 nt ramp region was markedly elevated (**Figure 6D**), accompanied by high RNA folding stability (**Figure S8L**) and a codon-usage skewed toward GC-rich codons for alanine, arginine, and leucine (**Figure 6E, G**). These amino acids are abundant in the amphipathic α-helices that form mitochondrial targeting signals ^26^, providing a structural rationale for the bias. Metagene analysis confirmed that high +60 GC content is globally associated with reduced ribosome occupancy in this region (**Figure S8P**), and this effect extends across the CDS, resulting in lower overall TE for hundreds of mitochondrial protein transcripts (**Figure 6H,I**),

Mitochondrial transcripts also tended to have extremely short 5′ UTRs (**Figure 6F**), which correlated with lower 5′ UTR ribosome occupancy (**Figure 6J**) and CDS translation in Liv^KO^ tissue (**Figure 6K**). UTRs shorter than ∼30 nt are thought to bypass canonical scanning and instead use alternative initiation routes ^27^, making them potentially dependent on specialised initiation factors. The enrichment of such short UTRs among mitochondrial mRNAs likely contributes to their selective impairment when BUD23 is lost. Very short 5′ UTRs also limit the possibility to harbour uORFs – which are effectively depleted from mitochondrial transcripts (**Figure S8M,N**).

To visualise these effects, we examined individual transcripts combining short 5′ UTRs with high +60 GC content – two features that synergise to confer strong BUD23 dependence (**Figure 6M-T**). *Bola3*, which encodes a mitochondrial [Fe-S] cluster assembly factor linked to multiple mitochondrial dysfunctions syndrome ^28^, has an 82% GC ramp region (encoding an amphipathic helix) and a 5′ UTR of <25 nt (**Figure 6I,K**). It showed a ∼4-fold reduction in CDS TE in Liv^KO^ (**Figure 6L**), with ribosome footprint read distribution revealing an absence in occupancy immediately after initiation and low coverage across the CDS (**Figure 6M**). Similar patterns were seen for *Cox6a1* (respiratory chain complex IV subunit) and *Ak3* (adenylate kinase essential for TCA cycle function) (**Figure 6N,O**). In other cases, such as *Uqcc2* (mitochondrial complex III assembly factor) (**Figure 6P**), TE reduction in Liv^KO^ (**Figure 6L**) occurred despite an average GC-content (61%; yet also encoding amphipathic helix), suggesting that the short UTR length alone can drive sensitivity. Similar conclusions could be drawn from non-mitochondrial metabolic factors such as *Agpat2* (**Figure 6Q**) that has a very short UTR and average GC content (68%). We also observed transcripts where *Bud23* loss selectively reduced uORF translation in the 5′ UTR, such as *Cisd3*, *Etfdh*, and *Igfbp4* (**Figure 6R-T**). Notably, the uORFs were located very close to the 5′ cap, a configuration associated with strong regulatory potential.

In summary, these analyses identify three sequence features – GC-rich ramp regions, short 5′ UTRs, and uORF content – as key determinants of BUD23-dependent translational initiation. Using differential protein abundance as a surrogate for Ribo-seq profiling, we find a similar association between both 5’ UTR length and GC-rich ramp region, and sensitivity to BUD23 loss within BAT (**Figure S6H**). The biases in these features in nuclear-encoded mitochondrial protein mRNAs explain their pronounced translational downregulation upon BUD23 loss.

### BUD23 expression is linked to human cardiometabolic and systemic traits

Given the metabolic and mitochondrial phenotype observed in *Bud23*-knockout mouse models, we next asked whether variation in *BUD23* expression in humans is linked with cardiometabolic health and related biomarkers. To do this, we applied Mendelian randomization (MR), a method that uses germline genetic variants as instrumental variables to estimate the causal effect of modulated gene expression on disease risk ^29,30^. Specifically, we used cis-eQTLs (expression quantitative trait loci), variants within the *BUD23* loci that were found to associate with *BUD23* expression in whole blood and assessed their consistent effects on major cardiometabolic outcomes, including coronary artery disease (CAD), obesity, metabolic-associated steatotic liver disease (MASLD), cirrhosis, and circulating biomarkers of lipid, glucose, and liver function.

Genetically increased *BUD23* expression in whole blood is inversely associated with coronary artery disease (CAD), metabolic-associated steatotic liver disease (MASLD), and liver enzymes, aspartate aminotransferase (AST), suggesting beneficial effects of increased BUD23 levels (**Figure 7A**). We additionally observed significant inverse association with hypertension, aligning with an association with lower systolic blood pressure (SBP) levels, as well as with reduced myocardial infarction (MI) risk. By contrast, increased *BUD23* expression was positively associated with body mass index (BMI) and obesity, indicating a potential trade-off between BUD23 effects on adiposity and cardiometabolic protection. Notably, effect estimates were concordant across both primary and replication GWAS sources, indicating the robustness of inferred causal effects to population-specific biases. Similar associations emerge from analyses of BUD23 obligate partner, TRMT112, and METTL5, another methyltransferase partner of TRMT112 which modifies 18S rRNA close to the decoding centre (**Figure S9A,B,D**).

**Figure 7.**
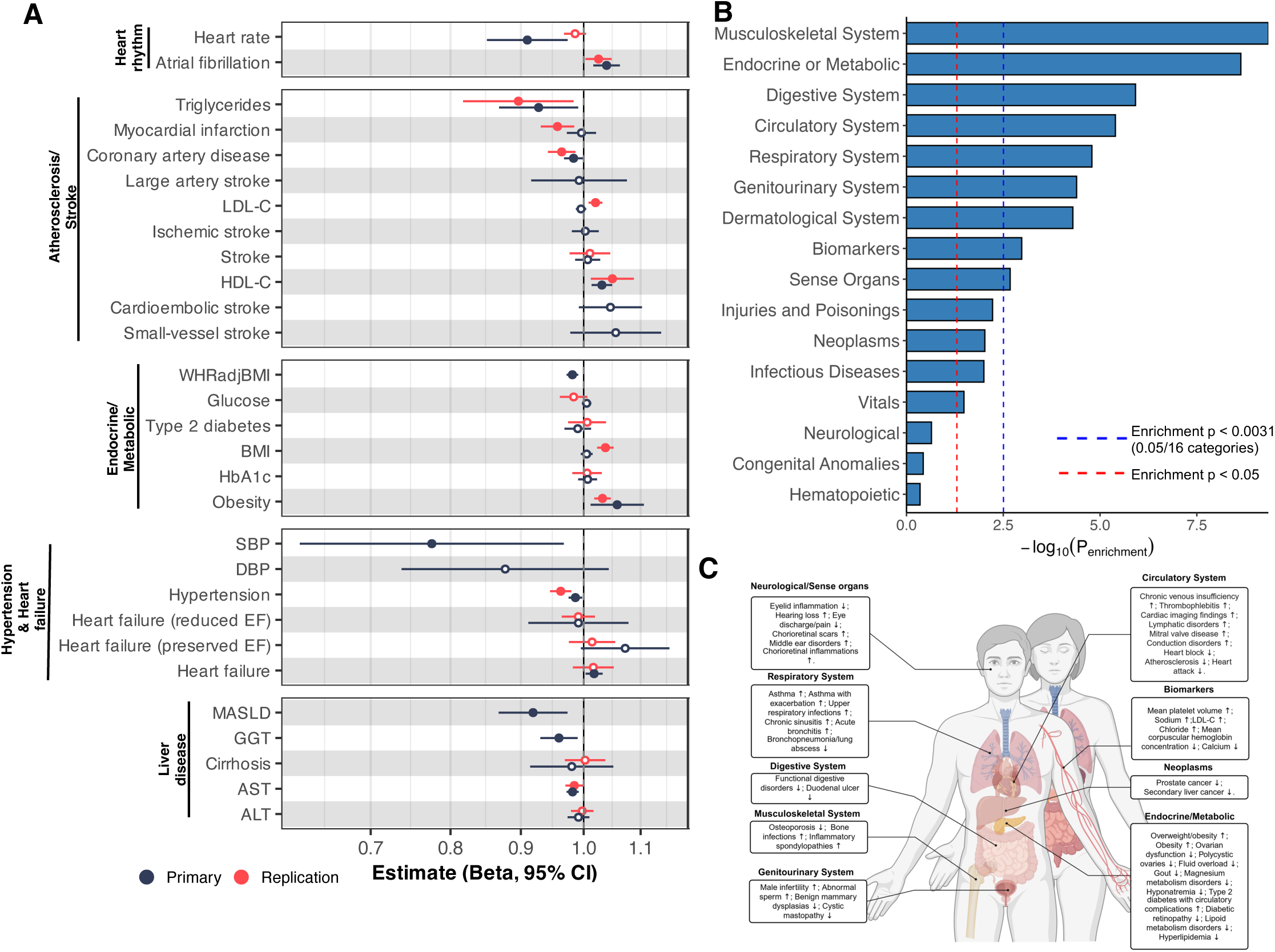
Human genetic evidence linking whole blood Bud23 expression to cardiometabolic disease outcomes. Mendelian Randomization (MR) analyses modelling the effect of genetically increased BUD23 expression in whole blood on human disease and biomarker traits. (**A**) Forest plots display MR estimates (Beta ± 95% CI) across cardiometabolic outcomes in both primary and replication GWAS datasets. Estimates reflect the direction and magnitude of association per unit increase in genetically predicted BUD23 expression. (**B**) Enrichment analysis summarizing phenome-wide MR associations across ∼1,200 traits in the Million Veteran Program (MVP), grouped by clinical domain. Bar heights represent enrichment significance, indicating domains with disproportionate representation of associated traits. (**C**) Homunculus schematic mapping representative traits that surpassed FDR correction (FDR < 0.05) within each disease category. Arrows denote the direction of association per increased BUD23 expression: upward arrows indicate positive associations with the outcome, while downward arrows indicate negative associations.

To test for tissue-specific effects, we repeated MR analyses using cis-eQTLs for *BUD23* expression in adipose (subcutaneous and visceral), skeletal muscle, heart (atrial appendage), and liver (**Figure S9C**). The direction and magnitude of effects in adipose tissues closely mirrored those in whole blood, particularly for obesity, lipids, and MASLD, suggesting shared metabolic pathways in these tissues. In contrast, muscle-derived *BUD23* instruments yielded directionally opposite effects, particularly for BMI and T2D, suggesting that BUD23 actions are tissue-specific.

To move beyond cardiometabolic phenotypes and capture the broader impact of *BUD23* on human health, we performed a phenome-wide MR scan across ∼1,200 traits from the Million Veteran Program (MVP) among participants of European ancestry ^31^. This analysis revealed significant causal associations between *BUD23* expression and traits related to metabolic, hepatic, and cardiovascular function, including liver fat, dyslipidemia, hypertension, and inflammatory biomarkers (**Figure 7B**). To contextualize these findings anatomically, we visualized significant phenotypes (FDR<0.05 per disease category) across disease domains using a homunculus-style plot (**Figure 7C**). The most prominent effects were observed in endocrine/metabolic, hepatic, and cardiopulmonary systems, reinforcing the hypothesis that BUD23 is a key regulator of systemic energy homeostasis.

Together with our mouse data, these results suggest that BUD23’s role in maintaining efficient translation of mitochondrial and metabolic transcripts extends to human physiology, influencing systemic energy balance and cardiometabolic disease risk.

## Discussion

The m^7^G modification is best known as the defining feature of the eukaryotic mRNA 5′ cap (m^7^GpppN) and is also found in tRNA, where it contributes to structural stability ^32^. In cytosolic ribosomes, methylation at G1639 is the only known – and evolutionarily conserved – m^7^G, installed by the 18S rRNA methyltransferase BUD23:TRTM112, which has no other known substrates ^8^. Despite its uniqueness, the molecular and physiological functions of this modification have remained largely unexplored. Here, we reveal that BUD23 action serves to modulate ribosomal translation of specific mRNA species. Those transcripts sensitive to BUD23 are enriched for mitochondrial factors and metabolic function providing ribosome-modification based control over cellular energy state.

### Structural basis of m^7^G1639 function in selective translation initiation

Our structural analyses reveal that m^7^G1639 occupies a central position in the initiating ribosome, directly contacting C41 in the anticodon stem of initiator tRNA, tRNA_i_^Met^, just downstream of the anticodon loop. The methyl group is crucial for positioning of the guanosine: it fills a hydrophobic pocket on the guanosine’s opposite face from the hydrogen-bonding edge contacting C41, and its positive charge enables cation-π stacking with Trp35 in RPS25. This provides greater stability than the π-π interaction associated with unmethylated G1639. A similar cation-π interaction between m^7^G and aromatic Trp side chains stabilises the mRNA cap within eIF4E ^33^.

That Trp35 is contributed by RPS25 is notable. This ribosomal protein flanks the mRNA exit tunnel, is present at sub-stoichiometric levels, and contributes to ribosome heterogeneity ^34^. It has been reported that RPS25-deficient ribosomes show reduced interaction with mitochondria-related transcripts ^34^. RPS25 is also required for initiating translation on highly structured viral IRES elements ^35,36^ and for repeat-associated non-AUG-initiated (RAN) translation of GC-rich repeats in FMR1 and C9orf72, associated with human neurodegenerative diseases ^37,38^. The structural proximity of m^7^G1639 and Trp35 of RPS25 at a key intermediate of 80S formation, and our finding that BUD23-sensitive transcripts are enriched for high GC content suggests an intersection between m^7^G1639 and RPS25-dependent mechanisms. This raises intriguing possibilities for combinatorial ribosomal heterogeneity involving these factors.

Although BUD23 has been considered a constitutive ribosome modification factor, emerging evidence suggests that rRNA modifications can vary across tissues and physiological states ^7^. Recent reports of variable methylation at the BUD23 target site ^39^ raises the possibility that m^7^G1639 installation is dynamically regulated, potentially allowing cells to adjust ribosome specialisation according to long-term metabolic demands. If so, BUD23 activity may be an important mechanism for regulatory coupling of energy state, translational output, and mitochondrial function. Indeed, recent evidence for specialised mitochondrially-located ribosomes as important for translation of nuclear-encoded mitochondrial mRNAs ^40^, may include location-specific heterogeneity in ribosome modification.

Related enzymes, such as the TRMT112 partner METTL5 ^41^, may also participate in a broader rRNA modification network with BUD23 that tunes ribosome function to metabolic demand. METTL5:TRMT112 install a nearby 18S modification m^6^A1832 that also modulates translation initiation ^42^. Loss of METTL5 results in metabolic phenotypes in mice ^41^ and our MR studies associate METTL5 expression with human cardiometabolic phenotypes.

### BUD23-dependent translational control converges on mitochondrial and lipid metabolism

Our findings reveal that the metabolic consequences of BUD23 loss in multiple tissues are rooted in a common mechanism: disruption of mitochondrial function and lipid metabolism. In WAT, *Bud23* deletion impairs lipid storage capacity *in vivo*, despite preserved adipocyte differentiation *in vitro*. This defect is accompanied by reduced expression of key lipogenic enzymes (e.g. *Fasn*, *Acaca*). These observations suggest that triglyceride synthesis and storage are critical targets of BUD23-directed rRNA modification. Mitochondrial dysfunction is likely to be central to the observed WAT phenotype, as mitochondrial activity in white adipocytes not only supports energy production but is also essential for lipid droplet formation and TG storage. Indeed, adipocytes rely heavily on mitochondria to maintain homeostasis, supporting processes including differentiation, lipogenesis, lipolysis, glucose metabolism, and adipokine secretion ^43^.

BAT is specialised for thermogenesis, with high mitochondrial density and pronounced expression of UCP1, which dissipates the proton gradient to generate heat ^44^. Mitochondrial fatty acid oxidation, TCA cycle activity, and electron transport are essential for this function ^45,46^. In *Bud23*-deficient mice, BAT thermogenesis appeared preserved, with body temperature and UCP1 expression unaltered under the conditions tested. This suggests that either mitochondrial changes were insufficient to compromise thermogenesis or compensatory pathways, such as proton leak via ANT1/2 ^47^, maintained heat production in the mice. Interestingly, “whitening” of BAT – characterised by lipid accumulation and a white adipocyte-like phenotype – was observed in Ad^KO^ but absent BAT^KO^ mice. This indicates that BAT whitening in Ad^KO^ mice is likely secondary to failure of WAT lipid storage, and highlights BAT’s role as a lipid sink and contributor to systemic lipid homeostasis ^48–50^.

Despite the distinct *Bud23*-deletion dependent phenotypes in WAT and BAT tissues, altered expression of mitochondrial factors, dysfunctional lipid storage and tissue loss were pronounced and common endpoints. This reinforces the importance of BUD23 activity in proper maintenance of mitochondrial dynamics, and highlights the now recognised dependence of adipocyte lipid handling (lipid droplet formation, rates of DNL/lipolysis, etc) on mitochondrial activity ^51–55^. Indeed, our transcriptomic and proteomic studies revealed several factors involved in inter-organelle coupling, lipid droplet formation/dynamics, and mitochondrial beta-oxidation which showed altered expression upon *Bud23* deletion.

In contrast to the pronounced phenotypes in adipose tissue, liver-specific *Bud23* deletion produced a milder metabolic phenotype – possibly due to the genetic targeting in adult mice and the relatively short time frame of our studies (2-4 weeks post-deletion). Nonetheless, clear alterations emerged: excessive hepatic lipid accumulation under normal and HFD feeding, and impaired fasting-induced β-hydroxybutyrate production. Both can be linked to mitochondrial dysfunction.

Ribosome profiling revealed that BUD23 loss shows significant selectivity towards impaired translation of nuclear-encoded mitochondrial transcripts, particularly those with short 5′ UTRs and high GC content downstream of the start codon. This GC bias is a predictable consequence of the amino acid composition of N-terminal mitochondrial targeting sequences. These features may hinder efficient initiation, making these mRNAs especially dependent on BUD23-modified ribosomes for effective translation. This model provides a unifying explanation for the mitochondrial defects observed across tissues.

The BUD23 dependent phenotypes we observed across tissues may reflect a combination of global effects – stemming from altered 40S maturation – and from transcript-selective effects on initiation mediated through m^7^G1639. However, the striking specificity for mitochondrial mRNAs argues for an active regulatory role for BUD23’s methyltransferase activity in shaping the translational landscape, rather than impacts on global ribosome assembly.

### BUD23 and nutrient-sensitive translational regulation

Our studies also uncovered an intriguing role for BUD23 in energy-sensitive translation. Under normal conditions, 5′ TOP mRNAs – encoding ribosomal proteins and translation factors – are tightly regulated by mTOR in response to nutrient availability. In *Bud23*-deficient livers, this regulation is lost, with 5′ TOP mRNA translation remains abnormally elevated during the light/fasting phase. This raises the possibility that BUD23-dependent rRNA modification is required for coupling nutrient sensing to translational control, possibly through effects on mTOR responsiveness. The consistently reduced insulin sensitivity observed *in vivo* and *ex vivo* in *Bud23*-deficient models further supports a defect in mTOR-mediated signalling. Two recent publications have provided structural and functional insights into how the TOP-binding repressive protein, LARP1, interacts with its substrate mRNA and with 40S subunits ^56,57^. It is intriguing to speculate that m^7^G1639 could be directly involved in the molecular mechanism of repression, although future experiments are required to explore this possibility.

### Relevance to human health

Finally, our study connects these mechanistic discoveries to human health, offering important insight into the role played by BUD23 and associated ribosome modification in health and disease states. BUD23 has been implicated in cancer and inflammation ^58–60^, and its gene lies within the critical deletion region of Williams-Beuren Syndrome (WBS), a multisystem disorder characterised by cardiovascular and metabolic abnormalities ^61^. Although multiple genes are deleted in WBS, our finding that *BUD23* is strongly associated with liver fat content (MASLD), body mass index, and obesity suggests it may meaningfully contribute to the metabolic features of this syndrome. Furthermore, protective associations for *METTL5* overlap with those for *BUD23*, reinforcing the concept that rRNA-modifying enzymes in the decoding centre contribute to metabolic homeostasis and are relevant to human cardiometabolic health and disease.

Together, our findings define BUD23 as a key regulator of ribosome specialization, selectively enhancing translation of mitochondrial mRNAs critical for energy metabolism. By linking nutrient sensing, translational control, and mitochondrial function, BUD23 emerges as a central node in the maintenance of metabolic homeostasis and a potential therapeutic target in cardiometabolic disease.

## Acknowledgment

We wish to thank and acknowledge support of the core facilities at the University of Manchester: Bioinformatics Core Facility, Genomic Technologies Core Facility, Biological Services Unit, and Histological Services Unit, aa well as technical assistance at the University of Oxford: Dr. Anne Clark, Electron Microscopy Facility (Dunn School), Amy Barret (biochemistry analysis), and the Target Discovery Institute Mass Spectrometry group (proteomics). We thank Drs. Eva Novoa and Jay Brito Querido for their valuable discussions and thoughtful comments, and Dr. Ana Domingos for providing access to the thermal imaging equipment. D.G. acknowledges funding by the University of Lausanne and by the Swiss National Science Foundation (SNSF grants 212423 and 10002692, and NCCR RNA & Disease, 205601). D.R acknowledges funding by NIHR Oxford Health Biomedical Research Centre (NIHR203316) and Medical Research Council (MRC; MR/W019000/1; MR/V034049/1). D.B acknowledges funding by the MRC (MR/P00279X/1) and Biotechnology and Biological Sciences Research Council (BBSRC; BB/V002651/1; BB/V002651/1)

## Author contributions

Conception: D.B., D.R., and D.G. Methodology: N.M-S, A.B., N.B., D.B.R, A.L, M.V., E.H., S.C., R.D., N.G., E.J., M.B., K.S., R.N., G.G., L.H. Original draft: N.M-S, D.B., D.R., and D.G. Revisions: N. M-S, A.B., D.B., D.R., and D.G. Supervision: D.B., D.R., and D.G.

## Material and Methods

### Animals

*Adipoq^Cre^Bud23^fl/fl^* (Ad^KO^), *UCP1^Cre^Bud23^fl/fl^* (UCP^KO^) and *Alb^CreERT2^Bud23^fl/fl^* (Liv^KO^) mice were generated by crossing *Adipoq^Cre^* (JAX laboratory, Strain #:028020), BAT^CRE^ (JAX laboratory, Strain #:024670) and Alb^CreERT2^ (given by Prof. Pierre Chambon, GIE-CERBM (IGBMC)) respectively, with *Bud23^fl/fl^* (generated at the University of Manchester (UK) ^11^. In all studies, CRE-negative littermates (Bud23^fl/fl^) were used as controls (designated: Ad^WT^; BAT^WT^; Liv^WT^). All mice were group-housed in 12:12 light/dark cycles, under controlled temperature (22±2°C) and humidity with ad libitum access to standard laboratory chow, unless stated otherwise. All studies used both male and female mice, unless otherwise stated. All experiments were carried out in accordance with the Animals (Scientific Procedures) Act 1986 (UK) under Home Office Project License PDC3CD59F (University of Oxford) and PP1136445 (University of Manchester) and approval from local ethical review bodies.

### *In vivo* treatments and studies

*Transgene induction*: Liv^WT^ and Liv^KO^ mice were treated daily with tamoxifen (Sigma, T5648) for 5 days (i.p., 0.1mg/day in sesame oil, Sigma S3547).

*High fat diet challenge*: Adult mice (10-13 weeks of age) were given ad libitum access to a high fat diet (60% energy from fat, DIO Rodent Purified Diet, IPS Ltd) for a period of 9-12 weeks.

*Metabolic phenotyping*: Body composition was assessed using an EchoMRI (Echo Medical Systems, E26-258-MT). Physiological (metabolic gas exchange) and behavioural (food and water intake, locomotor activity) rhythms were measured using the Phenomaster indirect calorimetry system (TSE Systems). Mice were individually housed and acclimatised for 24hr. with O2 consumption (VO2), CO2 production (VCO2) and energy expenditure (EE) recorded every 2 min for >72 hr. RER was derived from these measures (VCO2/VO2).

*Cold and thermoneutrality challenges*: For cold challenge, mice (∼13 weeks of age) were individually housed within a Phenomaster system, and following >2 days acclimatisation were exposed to an abrupt drop in ambient temperature to 4°C for 6 hours. For thermoneutrality studies, group housed mice were placed ∼29°C (+/-1°C) ambient temperature for 7 weeks.

*Insulin Tolerance Test*: Blood glucose was measured from tail blood using the Aviva Accuchek meter (Roche). For the insulin tolerance test, mice were fasted from ZT0, then injected with 0.75 IU/kg human recombinant insulin (I2643, Sigma-Aldrich) at ZT6 (time ‘0 min’).

*Thermal imaging & Body temperature:* BAT temperature of free moving Ad^WT^ and Ad^KO^ mice were measured using a thermal camera (Flir). Mice were shaved to expose the interscapular region >2 d prior to thermal imaging to avoid stress-induced BAT activity. Average temperature was calculated using Flir Tools software, where the average temperature was measured from a minimum of four images per mouse. Rectal temperature of unanaesthetised AD mice was measured using mouse rectal probe (RET-3, Type T Thermocouple, World Precision Instruments) connected to a BAT-12 Microprobe Thermometer (Physitemp Instruments). Recording of body temperature and activity was carried out via surgically implanted radiotelemetry devices (TA-F10, Data Sciences International). Following >10days recovery, body temperature was recorded every 5 min for >5 d.

*Labelled substrates study*. Male adult AD^WT^ / AD^KO^ animals were given deuterated water (^2^H_2_O) (CK Isotopes, Ibstock,UK) in the drinking water (25%vol/vol, ad libitum) for 48 hrs. After this period, they were fasted for 4 hours and then received an oral bolus of labelled glucose (D-glucose; U-13C6, Cambridge Isotope Laboratories, INC). Tissues were collected 2 hrs post-glucose administration.

### Primary cell culture

*Adipose tissue fractionation*: Brown adipose tissue (BAT) was collected and washed in PBS supplemented with amphotericin B from mice aged between 9-14 weeks old. The tissue was minced and digested in 1.5mg/ml collagenase H (25 min, shaking incubator 170rpm, 37°C). Collagenase solution was neutralised with DMEM (containing 20% FBS, 1% PS, 5µg/ml amphotericin B) and passed through a 100µm mesh filter. Cells were centrifuged (250g, RT 8 min) to separate the stromal vascular fraction (SVF) from the floating mature adipocyte layer. The SVF was collected and cultured until confluent. For BAT differentiation cells were plated at 50,000 cells/well, cultured for 3d prior to treatment with the differentiation cocktail (IBMX 0.5mM, indomethacin 0.125mM, dexamethasone 1µM, rosiglitazone 2.8µM, T3 1nM, insulin, 20nM) in DMEM (with 20% FBS, 1% PS). 5d post-plating, media was replaced with DMEM (with rosiglitazone 2.8µM, T3 1nM, insulin 20nM, 10% FBS, 1% PS). From 7d post-plating, media was replaced with Rosiglitazone (2.8µM) and Insulin (20nM) in DMEM + 10% FBS + 1% PS.

For gonadal white adipose tissue (gWAT) tissue was washed in PBS supplemented with amphotericin B, then minced and digested and washed as above. The gWAT SVF pellet was then collected and resuspended in DMEM (glutamax) + 20% FBS + 1% PS prior to isolation with the adipose tissue progenitor isolation kit (Miltenyi Biotec) according to manufacturer’s instructions. The purified progenitor population was cultured for 24 hrs in DMEM (20% FBS, 1% PS) before being replaced and maintained in DMEM (20% FBS only). Once confluent, cells were plated at 50,000 cells/well. Differentiation was triggered 3d later using differentiation cocktail (IBMX 0.5mM, Dexamethasone 1µM, Rosiglitazone 4µg/ml, insulin 5µg/ml in DMEM, 20% FBS). On day 6 and day 8 post-plating, media was replaced with DMEM (20% FBS, 5µg/ml insulin).

*Oil Red O and BODIPY/Hoechst staining.* Differentiated cells were stained using Hoechst 33342, Trihydrochloride, Trihydrate and BODIPY 493/503 (4,4-Difluoro-1,3,5,7,8-Pentamethyl-4-Bora-3a,4a-Diaza-s-Indacene; Fisher Scientific UK Ltd.) at 1:2000. Images were taken using the EVOS™ M5000 Imaging system. For Oil Red O staining (ORO), cells were washed in PBS and fixed in 10% Formalin (Merck Life Science UK Limited) for 30 mins. Cells were rinsed with deionised water and incubated with 60% isopropanol for 5 min before being stained with ORO solution for 15 minutes. Cells were rinsed with deionised water before being imaged on the EVOS™ M5000 Imaging system. Following imaging, the ORO was quantified by the addition of 100% isopropanol for 5 minutes. The absorbance of the extracted ORO solution was measured at 492nm on a plate reader.

*Mature adipocyte (MA) studies*. Following a method adapted from Rocha et al. ^62^, gWAT was collected from adult mice and washed in Hanks’ Balanced Salt Solution (Sigma). Tissue was minced and digested in 1 mg/ml collagenase (Collagenase H, Sigma) for 30 min in a shaking incubator at 170 rpm, 37°C. The sample was then centrifuged at 1000 rpm for 5 min at 4°C. MA (floating layer) was collected separately, lysed in TRIzol Reagent (Invitrogen), and stored at −80°C before proceeding to RNA extraction. To remove excess lipid from MA fractions, samples were centrifuged (full speed, 5 min, room temperature). RNA extraction was then carried out as per the manufacturer’s TRIzol protocol, up to the stage of removing the isopropranol phase, which was transferred to Reliaprep columns (Promega) for on-column DNase treatment, clean-up, and elution as per manufacturer’s protocol.

*Cell culture RNA extraction.* RNA was extracted from cells using the ReliaPrep™ RNA cell mini prep system (Promega UK Ltd.) following the manufacturer’s instructions. Samples were DNAse treated during the extraction procedure. RNA was eluted from the column in a final volume of 15µL and RNA concentration and purity was assessed using a NanoDrop 2000c UV/IV Spectrophotometer.

### Transcriptional Analyses

*RNA extraction (tissue)*. Total RNA was isolated by using Trizol Reagent (Invitrogen; Carlsbad, CA, USA) according to the manufacturer’s protocol (RNA was precipitated with chloroform and isopropanol, washed with 75% ethanol, and finally dissolved in RNase-free water). RNA concentration and purity was assessed using a NanoDrop 2000c UV/IV Spectrophotometer. Before the retrotranscription, RNA was DNAse treated using RQ1 DNAse (Promega) following manufacturer’s protocol.

*RT-PCR.* Samples were DNase-treated (RQ1 RNase-Free DNase, Promega, Madison, WI) prior to cDNA conversion High Capacity RNA-to-cDNA kit (Applied Biosystems). qPCR was performed using a GoTaq qPCR Master Mix (Promega, Madison, WI) and primers listed in Appendix Adipocyte NR1D1 dictates adipose tissue expansion during obesity using a Step One Plus (Applied Biosystems) qPCR machine. Relative quantities of gene expression were determined using the [delta][delta] Ct method and normalised with the use of a geometric mean of the housekeeping genes Hprt, Ppib, and Actb. The fold difference of expression was calculated relative to the values of control groups.

*RNA-Sequencing.* gWAT was collected from adult male mice and flash-frozen. Total RNA was extracted and DNase-treated as described above. Biological replicates were taken forward individually to library preparation and sequencing. For library preparation, total RNA was submitted to the University of Manchester Genomic Technologies Core Facility (GTCF). Quality and integrity of the RNA samples were assessed using a 2200 TapeStation (Agilent Technologies) and then libraries generated using the TruSeq Stranded mRNA assay (Illumina, Inc) according to the manufacturer’s protocol. Briefly, total RNA (0.1–4 μg) was used as input material from which polyadenylated mRNA was purified using poly-T, oligo-attached, magnetic beads. The mRNA was then fragmented using divalent cations under elevated temperature and then reverse-transcribed into first strand cDNA using random primers. Second strand cDNA was then synthesised using DNA Polymerase I and RNase H. Following a single ‘A’ base addition, adapters were ligated to the cDNA fragments, and the products then purified and enriched by PCR to create the final cDNA library. Adapter indices were used to multiplex libraries, which were pooled prior to cluster generation using a cBot instrument. The loaded flow cell was then paired-end sequenced (76 + 76 cycles, plus indices) on an Illumina HiSeq4000 instrument. Finally, the output data was demultiplexed (allowing one mismatch) and BCL-to-Fastq conversion performed using Illumina’s bcl2fastq software, version 2.17.1.14.

RNA was extracted from BAT using the SV Total RNA Isolation System (Promega) according to manufacturer’s instructions. RNA yield was quantified by TapeStation (Agilent), to ensure it was of sufficient quality for sequencing. Library preparation and sequencing for the Illumina HiSeq 4000 platform were performed by Novogene. Raw FASTQ files were processed through a standard pipeline by Novogene to generate a list of counts. Gene lists were analysed for differential expression using a combination of techniques, including edgeR ^63^.

### Genomic DNA extraction and Mitochondria Copy Number analysis

Genomic DNA (gDNA) was extracted using phenol-chloroform. Tissue was homogenized in TE buffer and incubate at 55°C for 3 hours with proteinase K and 20% SDS. After that, 1 volume of phenol:chloroform:isoamyl alcohol (25:24:1; Thermo Fisher) per sample was added and vortexed for 20 seconds. Samples were transfered into phase lock tubes (Qiagen) prior to centrifugation at 16,000g for 5 mins. The aqueous phase was carefully removed and transferred to a fresh tube. 1 μl GlycoBlue, half a volume of 7.5M ammonium acetate, and 2.5 volumes of 100% ethanol were added. The sample was kept at –80°C for one hour to facilitate DNA precipitation. The sample was then centrifuged at 16,000g at 4°C for 30 mins to pellet the DNA. The pellet was washed in ethanol once and left to air dry. Purified genomic DNA was resuspended in nuclease-free water.

Mitochondria Copy Number was analysed by qPCR by measuring the mitochondrial genome gen mt-ND1 and normalizing the values by genomic GAPDH.

### Mitochondrial function

Liver was isolated and transferred to isolation buffer (in mM: sucrose 200, MOPS 10, EGTA 1, pH 7.4 adjusted with KOH). The tissue was minced, homogenised with three passes of a glass tissue homogeniser and homogenate centrifuged at 7000 x g (all centrifugation steps for 10 minutes at 4 °C). Excess fat from the centrifugation tube was removed and the pellet resuspended in isolation buffer before further centrifugation at 600 x g. The supernatant was filtered through gauze before centrifugation at 7000 x g. The resultant pellet was resuspended in isolation buffer (14.8 ± 2.8 mg/ml) and kept on ice.

Mitochondria (173 ± 22 µg) were loaded into an Oroboros O2k high resolution respirometry system (Oroboros Instruments, Innsbruck, Austria) containing MIR05 buffer (0.5 mM ethylene glycol-bis(ß-aminoethyl ether)-N,N,N’,N’-tetraacetic acid, 3 mM MgC_l2_, 60 mM K-MES, 10 mM KH_2_PO_4_, 20 mM 4-(2-hydroxyethyl)-1-piperazineethanesulfonic acid, 110 mM sucrose, 0.1% bovine serum albumin (BSA), pH 7.1 adjusted with KOH) for measurement of mitochondrial respiration rate. All measurements of respiration rates were carried out at 30°C. Oxygen electrodes were calibrated daily with air-saturated respiration solution.

OXPHOS and LEAK were measured in the presence of Complex I substrates pyruvate (6.25 mM), glutamate (10 mM) and malate (2 mM). The protocol used to measure these parameters was adapted from ^64^. Briefly, pyruvate (6.25 mM), malate (2 mM) and glutamate (10 mM) are added as carbon substrates and to spark the citric acid cycle. Under these conditions, mitochondria are in LEAK respiration (State II) with CI substrates in the absence of adenylates. OXPHOS with CI substrates (State III) was achieved through addition of saturating levels of ADP (125 µM). State IV was then measured once ADP had been depleted, and the RCR was calculated by State III / State IV. Results were normalised to protein content determined using the Bradford method.

### Ribosome profiling

Livers from Liv^KO^ and age-matched Liv^WT^ animals (both groups tamoxifen-treated) were collected at diurnal Zeitgeber timepoints ZT6 and ZT18 (3 male mice per genotype and timepoint) and flash-frozen in liquid nitrogen. Using ∼200 mg of frozen sample per liver, tissue lysates were prepared and ribosome footprints were generated (RNase I) and purified, all according to previously described protocols as in Janich et al ^19^. An aliquot of the same lysate as for footprint generation was used to purify matching total RNA preparations, of which 1 μg was chemically fragmented for RNA-seq library preparation, also as described in ^19^. Size-selected, fragmented RNA and footprint samples were then subjected to library preparation protocols identical to those used in our previous studies (e.g. ^65^. In this protocol, sample barcodes and unique molecular identifiers (UMIs) are included in the initial adaptors ligated to the RNA molecules (similar to ^66^), allowing for multiplexing at an early stage, before rRNA depletion (6 samples were multiplexed for each of the final four libraries). The amplification of the libraries was carried out using i5 and i7 indexed primers (12 PCR cycles). The libraries were sequenced on a NovaSeq6000 (Illumina).

*Ribosome profiling data mapping*. Read mapping was performed essentially following our published protocol ^67^. Briefly, reads were trimmed from adapter sequence using cutadapt (version: 3.5; options: –-match-read-wildcards –-overlap 8 –-discard-untrimmed –-minimum-length 30) and quality filtered using fastx_toolkit (version: 0.0.14; options: –Q33 –q 30 –p 90). UMIs were extracted from each read with UMItools (version: 1.0.0git; options: extract –-extract-method string –-bc-pattern NNNNNNNNCCCCC –-3prime –-filter-cell-barcode –error-correct-cell). Then, reads where size-selected for monosome footprints (size 26 to 35). Subsequently, to estimate rRNA and tRNA contamination, reads were mapped to human and mouse rRNA and mouse tRNA databases using bowtie2 (version: 2.3.5; options: –p 2 –L 15 –k 20 –-trim5 2). Reads that failed to map to these, were then mapped to the mouse transcript database (Ensembl database v. 100). Barcode demultiplexing was carried out with UMItools (options: group –-method=directional –-per-cell –read-length) and deduplication with an in-house script. For each gene, only one transcript isoform was considered, namely the primary isoform based on classification by the APPRIS database ^68^. In case several transcript isoforms were annotated as primary by APPRIS, the one with the longest coding region was selected. For Ribo-Seq reads, the ribosome A-site was assumed to cover nucleotide positions 15-17, and Ribo-Seq reads were counted for each gene that overlapped with their A-site the coding region. For RNA-Seq, all reads were counted that aligned to a gene.

*Data and code availability.* For liver ribo-seq and matching RNA-seq data, raw sequencing data files will be deposited in NCBI’s Gene Expression Omnibus (GEO) archive. The scripts for data analysis will be abailable from https://github.com/gatfieldlab/.

*PCA analysis.* PCA analysis was done in python3 using the scikit-learn package ^69^: Scikit-learn: Machine Learning in Python ^70^ considering genes with any reads in at least 10 out of 12 samples. Read counts were normalized by size factors obtained by DESeq2 ^71^. Translation efficiencies were calculated by dividing normalized Ribo-Seq read counts by normalized RNA-Seq read counts for genes with any RNA-Seq reads.

*Differential expression and ribosome occupancy analysis.* Differential expression and ribosome occupancy analysis was done using DESeq2 ^71^ considering RNA-Seq and Ribo-Seq read counts, respectively. Differential translation efficiency analysis was done in python3 considering genes with reads in all RNA-Seq and Ribo-Seq samples. An independent T-test was used to compare translation efficiencies in Bud23-KO with wildtype samples, and p-values were FDR corrected. Genes with corrected p-value < 0.05 and absolute log2-fold change > 1.5 were considered significantly differentially translated.

*GO analysis and gene sets.* GO enrichment analysis was done in python3 using gene ontology gene sets downloaded from MSigDB (https://www.gsea-msigdb.org/gsea/index.jsp) with at least 20 genes. For each gene set, the proportion of genes contained in the set among significantly changed genes (in KO versus WT, based on RNA-Seq, Ribo-Seq or TE) was compared with the proportion for all genes using Fisher’s exact test, and *p*-values were FDR-corrected. If all genes of a smaller gene set were contained in a larger gene set and both gene sets were significantly depleted or enriched, only the larger gene set was retained. Selected representative gene sets are shown in **Figure 5D**. In **Figure 5E**, the TE log2 fold changes of genes in a GO set were compared with those of all genes using a Wilcoxon rank-sum test, and *p*-values were FDR-corrected.

In Figure 6, the ribosome gene set is from the GO cellular component (CC) ontology, excluding mitochondrial ribosomal genes (Mrp genes), and the mitochondrial gene set is from MitoCarta3.0 (downloaded from https://personal.broadinstitute.org/scalvo/MitoCarta3.0/Mouse.MitoCarta3.0.xls)

*Codon dwell time estimation.* Codon dwell times (DTs) were estimated using the RiboDT pipeline ^72^, based on Ribo-Seq reads. In **Figure S6F**, DTs when assuming an interaction between P– and A-sites are shown, as well as the difference between both (KO – WT).

*Meta-profile of ribosome occupancy around CDS starts and ends.* For each Ribo-Seq read, the A-site was assigned to start at the nucleotide at positions 15. The number of A-sites of ribosomes were counted at each nucleotide in a region from 100 nucleotides upstream to 200 nucleotides downstream of annotated start codons and in a region from 200 nucleotides upstream to 100 nucleotides downstream of annotated stop codons. For each gene, A-site counts of different samples were normalized by their sizefactors (obtained from DESeq2), summed over all Liv^KO^ or all Liv^WT^ samples, and then z-scored relative to the profile counts for wildtype. Genes with A-site counts at more than 25 positions were considered. **Figure 5G** shows the average (line) and standard error (shaded area around line) for Liv^KO^ and Liv^WT^ samples. A one-sample t-test was used to test, at each position, if the distribution of differences of z-scored A-site counts between KO and wildtype for all genes was significantly smaller or larger than 0 (i.e. no difference). The p-values for all positions were FDR corrected and considered significant when < 0.05.

*Sequence properties of transcripts and their correlation with changes in translation efficiency (TE) for specific gene regions.* We quantified several sequence properties of transcripts and examined their correlation with TE fold-changes in specific gene regions: in 5’ UTRs, in coding regions, and in first or last 60 nucleotides of coding regions. We considered the following transcript sequence properties: distance of start codon from mRNA 5’ end (5’ UTR length), number of upstream open reading frames (uORFs) in 5’ UTRs starting with a AUG start codons, an in-frame stop codons, and ribosome A-site occupancy, number of uORFs starting with a (C/G/U)UG start codon, Kozak score, codon optimality score (for first 20 codons), GC content for different regions (5’ UTRs, 60 nucleotides upstream or downstream of start codons or both), absolute minimum free energy (MAF) for folded RNA within 60 nucleotides upstream or downstream of start codons or 120 nucleotides surrounding start codons).

As 5’ UTR length we considered two measures: the distance between the annotated translation start codon and the annotated mRNA 5’ end, and the distance between the annotated translation start codon and the start of transcription as indicated by CAGE (Cap analysis of gene expression) data obtained from mouse liver by the Fantom Consortium (data downloaded from https://fantom.gsc.riken.jp/5/datafiles/latest/basic/). In particular, we assigned the transcription start site to the position with the maximum number of CAGE reads (at least 10 reads) within the first annotated exon and 500 nucleotides upstream of it. CAGE data for adipocyte analyses was treated analogously (using datasets mesenchymal stem cell differentiation to adipocytes, day06, biol_rep 1 to 3).

The codon bias score was determined by summing up the relative abundances of codons, corresponding to each codon within the first 20 codons, where the relative codon abundances were calculated from codons in expressed genes and weighted by the gene expression level in wildtype samples.

Spearman correlations between transcript sequence properties and TE fold-changes were calculated for genes with more than 1 normalized RNA-Seq read in KO and WT, more than 10 normalized Ribo-Seq reads in KO or WT, and a ratio of normalized Ribo-Seq to RNA-Seq reads larger than 0.5 for KO or WT. As several sequence properties correlated across transcripts, we selected representative sequence properties.

*Linear regression model to explain TE FCs with sequence properties.* Linear regression was used to select gene sequence properties that together contribute to explaining the variance in TE FC and to quantify the total fraction of the variance explained by these combined gene sequence properties. To select contributing sequence properties, we performed a forward feature selection procedure, where in each step the sequence property maximizing the model correlation with TE FC was added. We used a five-fold cross-validation to evaluate the model performances, with regression coefficients being estimated on 80% of the data and correlations of the parametrized models with TE FCs being evaluated on the remaining 20% of the data. Splitting of the data was random and repeated 50 times. A new gene sequence property was added to the model if it increased the explained variance of TE FCs by at least 1% compared to the explained variance by the previous model.

### Stable isotope analysis/Gas chromatography

Total lipids were extracted from tissue homogenates using the Folch method (chloroform:methanol (2:1; v/v) ^73^ and separated by solid-phase extraction (SPE) into lipid fractions and fatty acid methyl esters (FAMEs) as previously described ^74^. Deuterium incorporation from ^2^H_2_O in plasma water was measured using a Finnigan GasBench II (Thermo Fisher Scientific, Paisley, UK). Tissue palmitate ^13^C (from U-^13^C_6_ glucose) and ^2^H (from ^2^H_2_O, heavy water) enrichment was determined by GC-mass spectrometry (GC-MS) (Agilent Technologies; CA, USA) by monitoring ions with mass-to-charge ratios (m/z) as follow: 270 (M+0), 271 (M+1) and 272 (M+2).

### Western blotting

Total protein was isolated from gWAT and liver using protein extraction RIPA supplemented with protease and phosphatase inhibitors in a tissue homogenizer. Protein quantification was performed following the Bradford method. Protein lysates were subjected to SDS-PAGE, electrotransferred and blocked with milk/BSA. Primary antibodies were incubated overnight at 4C. Fluorescence secondary antibodies were incubated 1 hour at room temperature and imagines were taken using Biorad Gel Doc system. Band signal was quantified by densitometry using ImageJ 1.33 software, values were expressed in relation to b-actin/ total protein ponceau. Representative images for all proteins are shown.

### Histology

Haematoxylin and Eosin staining was carried out on paraffin embedded sections using the Leica ST5010 Autostainer XL. Images were acquired on a 3D-Histech Pannoramic-250 microscope slide-scanner using a 20x/ 0.80 Plan Apochromat objective (Zeiss). Snapshots of the slide-scans were taken using the SlideViewer software (3D-Histech). Further imaging was also carried out on the Zeiss AX10. Adipocyte diameter was quantified using ImageJ and the Adiposoft plugin version 1.16.

### Proteomics – Mass Spectrometry

Adipose tissue was homogenised using a TissueRuptor II (Qiagen) in 1x TBS. SDS was added to a final concentration of 4%, and the lysate was immediately boiled at 95°C for 10 minutes. Homogenates were then processed for an in-solution protein digest using a modified methanol/chloroform protein extraction method. Briefly, proteins were reduced with 5mM DTT (60 min, room temperature, with agitation) followed by alkyation with 20mM iodoacetamide (60 mins, dark). Lysates were vortexed in 60% methanol 15% chloroform before centrifugation at max speed in a bench-top centrifuge for 1 minute. Aqueous phase was then removed. The organic phase was then washed with methanol prior to centrifugation for 2 mins at max speed. The resulting pellet was resuspended in 6M urea, which was then diluted down to <1M prior to protein quantification (Bradford assay). Samples were digested overnight at 37°C with agitation (1,500 rpm, ThermoMixer) with trypsin (1:50 enzyme-to-protein ratio). Peptides were acidified using TFA and then desalted during Pierce C18 tips (Thermo Scientific) per manufacturer’s instructions. Eluted peptides were dried in a SpeedVac and resuspended in 0.1% formic acid for mass spectrometry analysis.

Peptides were analysed by nano-UPLC-MS/MS using a Dionex Ultimate 3000 coupled to an Orbitrap Fusion Lumos (Thermo Scientific) using a 75 µm x 500 mm C18 EASY-Spray Columns with 2 µm particles (Thermo Scientific) at a flow rate of 250 nL/min operated in trap and elute mode using a 60 minute linear gradient from 2% buffer B to 35% buffer B (buffer A: 5% DMSO, 0.1% formic acid in water; buffer B: 5% DMSO, 0.1% formic acid in acetonitrile). MS1 scans were acquired positive mode in the Orbitrap using a scan range of 400-1500 m/z, resolution of 120K, AGC target of 4 × 105, and maximum injection time set to auto. MS/MS scans were acquired in the ion trap using rapid scan mode. Precursors were selected for HCD fragmentation with a normalised collision energy of 28% using a 1 second cycle time (charge states 2-7, minimum intensity threshold 5,000, dynamic exclusion window of 60 seconds).

Raw data files were searched against the UniProtKB mouse database (retrieved 01/08/2021) using MaxQuant v1.6.0.16. Enzyme was set to trypsin with up to 2 missed cleavages, and match between runs was enabled. Methionine oxidation and protein N-terminal acetylation were set as variable modifications, and cysteine carbamidomethylation was set a fixed modification. All other settings were left as default. Output data was further processed using R statistical software (R Core Team, 2017; Wickham, 2009) and Perseus version 1.6.0.7 (Tyanova et al., 2016). Proteins were filtered to remove any entries that were flagged as potential contaminants, part of the decoy reverse database, identified by site only, or belonging to the ‘blood microparticle’ GO term. Any protein that was only identified in less than half of the total number of samples was removed. All label-free quantification values were log(2) transformed. Missing values were imputed using a normal distribution (width = 0.3, downshift = 0.18 standard deviations) ^75^.

### Blood biochemistry

Blood serum was acquired via cardiac puncture. Blood was allowed to clot at room temperature for 30 mins, before centrifugation at 2,000g for 10 mins. Serum was collected and the pellet discarded. Metabolites (including glucose, glycerol, NEFA, TG, lactate, cholesterol, 3-OHB, CRP, Urea, HDL) were assessed via ILab 650 Automatic Biochemistry Analyzer Clinical Chemistry System.

*Adiponectin ELISA*. Adiponectin in cell culture supernatant was measured using the mouse Adiponectin/Acrp30 DuoSet ELISA (Bio-Techne Ltd.) according to manufacturer’s instructions. Cell culture supernatant was diluted 1:500 to fall within the range of the assay.

*Bio-Plex*. Serum samples were analysed using the Bio-Plex Pro Mouse Diabetes 8-Plex assay kit (171F7001M, Bio-Rad Laboratories Ltd) and the Bio-Plex 200 system (Bio-Rad Laboratories Ltd.). The assay was run according to the manufacturer’s instructions and at the recommended dilution for serum samples.

### Triglyceride assay

Triglyceride in tissue lysates was measured using a Cayman’s Triglyceride calorimetric assay (CAY700190-96 wells, Cambridge Bioscience Ltd.) according to the manufacturer’s instructions. Lysates were prepared from ∼100mg pieces of liver homogenised in lysing matrix D tubes (MP Biomedicals) with NP40 substitute assay reagent supplemented with a cOmplete mini EDTA protease inhibitor tablet (Merck Life Science UK Limited). Lysates were diluted 1:3 to fall within the range of the assay.

### Electron Microscopy

Livers were dissected from adult mice, after PBS perfusion and immersed in fixative solution (2.5% glut/4% formaldehyde) for 1 hr at room temperature and then kept at 4C until processing. Fixed samples were processed with microwave assistance using a Leica AMW according to the following steps. Samples were washed with buffer (0.1 M sodium cacodylate buffer pH 7.2), stained with 1% osmium tetroxide and 1.5% potassium ferrocyanide in buffer, then rinsed with MilliQ water. Samples were further stained with 2% uranyl acetate in water, rinsed again with MilliQ water, then dehydrated through an ethanol series (30%, 50%, 70%, 90%, 95% and absolute ethanol). Samples were then infiltrated with 25%, 50%. 75% and finally 100% low viscosity resin (TAAB) in ethanol. Samples were removed from the microwave processor and submerged in fresh 100% resin, then placed on a rotator overnight. Samples were incubated on the rotator for the following 4 days, with changes into fresh resin twice per day. The samples were embedded on the afternoon of the final day and polymerised at 60°C for 48 hours. Sections of 90 nm were cut from the resin blocks using a Leica UC7 Ultramicrotome and collected onto 3 mm copper grids. The sections were then post-stained with lead citrate and imaged using a JEOL Flash 120kV TEM equipped with a Gatan Rio camera. Analysis was done using Image J/Fiji. Mitochondrial area was measured using manual segmentation of the outer membrane. For cristae density, images were inverted and, for each mitochondrion, a segmented line was drawn along the major axis and the corresponding intensity profile was extracted. Cristae were identified as local intensity maxima, and the number of detected peaks was divided by the mitochondrial area (µm²) to calculate cristae density (peaks/µm²), as previously described ^76^.

### Structural models

For structural models in **Figure 5I** and **Figure S7**, PDB entries 8pj1-5 were loaded in Swiss-PdbViewer ^77^ and the m^7^G1639 methyl group was added by superposing the guanine aromatic ring of pdb entry 5H3T chain A:MGT801 onto PDB entries 8pj1-5 G1639 guanine aromatic rings. The molecular surface of 18S rRNA (chain A) was computed excluding G1639 to show the space available to perfectly accommodate m^7^G1639. Chains e (40S ribosomal protein S25/ eS25), f (40S ribosomal protein S18 / uS13), w (initiator Met-tRNA) and 7 (mRNA) were drawn in purple, teal, dark green, yellow ribbon, respectively.

### Human genetic analysis

*Construction of Genetic Instruments.* To model genetically predicted expression of BUD23, TRMT112, and METTL5, we used significant cis-eQTLs (P < 5×10^-8^) located within ±500 kb of each gene. Independent variants were selected using LD clumping at R² < 0.1 based on European 1000 Genomes reference data ^78^. Whole blood served as the primary tissue, using eQTL data from the eQTLGen consortium ^79^. For tissue-specific sensitivity analyses, we used GTEx v8 data ^80^ for skeletal muscle, adipose visceral, and heart (atrial appendage), and subcutaneous adipose ^81^. Tissues were selected based on their relevance to metabolic disease and data availability.

*Outcome Selection.* Primary outcomes were selected to capture the breadth of cardiometabolic risk, spanning cardiovascular, hepatic, lipid, glucose, and adiposity domains. These included coronary artery disease (CAD), myocardial infarction (MI), hypertension, systolic blood pressure (SBP), type 2 diabetes (T2D), obesity, and body mass index (BMI). Additional outcomes included canonical biomarkers of hepatic (ALT, AST, liver fat), lipid (LDL-C, HDL-C, triglycerides), and glucose metabolism (HbA1c), as well as inflammation (C-reactive protein, CRP). These traits were selected a priori based on known metabolic relevance and visualized in the primary MR forest plot (see Figure 7A). All outcome GWAS were derived from large-scale European ancestry meta-analyses or biobank-based cohorts. For replication, we used summary statistics from the Million Veteran Program (MVP) ^31^, including a second CAD GWAS ^82^ and corresponding endpoints. Outcomes were harmonized to ensure consistent effect allele alignment with eQTL instruments.

*Phenome-Wide Mendelian Randomization and Enrichment.* To explore broader phenotypic consequences of gene perturbation, we conducted a phenome-wide MR (PheWAS-MR) scan of BUD23 across 1,022 traits in the MVP. Traits were filtered to exclude hierarchical or aggregate groupings (e.g., “any cardiovascular disorder”), allowing more granular resolution. Traits were classified into clinical domains using MVP’s predefined disease categories (e.g., cardiovascular, hepatic, endocrine/metabolic, inflammatory). We then tested for enrichment within each domain by assessing whether the distribution of MR P-values deviated significantly from the expected null (uniform) distribution. This enrichment-based prioritization approach, adapted from Schmidt et al. ^83^, allows for identification of directional or pleiotropic signatures across phenotypic systems, even when individual traits may not surpass genome-wide significance thresholds. This method balances the need for power (i.e., false negative control) against the risk of false positives and is particularly relevant when using MR to identify drug repurposing opportunities or detect early safety signals.

*Quantification and statistical analysis.* Two-sample MR was performed using the inverse-variance weighted (IVW) method as the primary estimator. For exposures with only one SNP instrument, the Wald ratio method was used. All results were aligned to the effect of increased gene expression and are reported as β-coefficients (or log odds for binary traits). Nominal significance was defined as P < 0.05 for the main cardiometabolic outcomes.

For phenome-wide MR, false discovery rate (FDR) correction was applied within each disease domain. For domain-level enrichment, deviation from the null distribution of P-values was tested using Kolmogorov-Smirnov or similar non-parametric methods, as in prior enrichment MR frameworks. All statistical analyses were conducted in R (v4.2.2) using the TwoSampleMR package ^84^. P-value distribution plots and enrichment histograms were generated using custom R and Python scripts.

### Data analysis and statistics

Data are presented as mean +/-standard error of the mean (SEM). Significance was defined as p <0.05 and the significance level is indicated in figure legends. Statistical significance was determined by Student t test (when two groups were compared), and two-way analysis of variance (ANOVA) when more than two groups were compared (2 factors) followed by Šídák post hoc test. Data analysis and graphing were performed in GraphPad Prism, R, Phyton and Adobe Illustrator.

---

**Figure S1.**
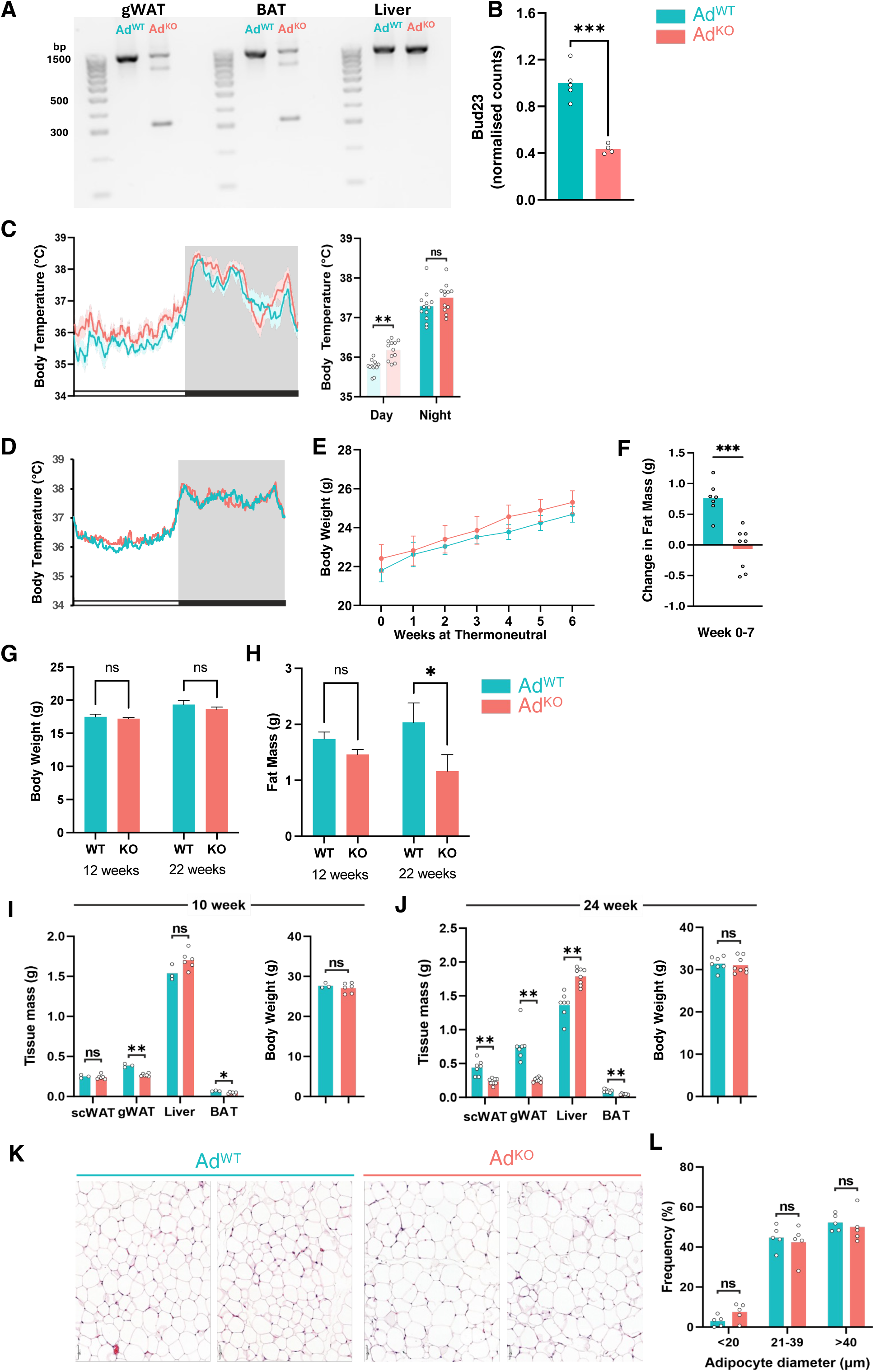
Loss of Bud23 function drives profound metabolic phenotype. (**A**) Representative agarose gel image confirming recombination at *Bud23* loci in gonadal white adipose tissue (gWAT) and brown adipose tissue (BAT). Liver has been included for comparison, serving as a negative control and highlighting tissue selectivity. Band at ∼350bp in gWAT and BAT of Ad^KO^ animals indicates recombination. (**B**) *Bud23* expression in gWAT of Ad^WT^ and Ad^KO^ mice. (**C**) Body temperature prolife and day/night average for Ad^WT^ and Ad^KO^ mice (n=12). (**D-F**) Body temperature (**D**), body weight (**E**), and change in fat mass (**F**) of Ad^WT^ and Ad^KO^ mice housed at thermoneutrality (28°C) for 7 wks (n=7-8). (**G**, **H**) Body weight (**G**) and fat mass (**H**) of female Ad^WT^ or Ad^KO^ 12 and 24 wks of age. (**I, J**) Tissue mass and body weight of male Ad^KO^ mice and their littermate controls at 10 and 24 wks of age. (**K, L**) H&E representative images and adipocyte size profiles (diameter in μm) in gonadal gWAT of Ad^KO^ and Ad^WT^. n reflects biological replicates (mice) in all panels. Mean values are plotted (with ± SEM where appropriate). Statistical significance was tested by unpaired two-tailed Student’s t-test (B, D, H, I) or Two-way ANOVA with Holm-Šídák post hoc test (C, K). *p<0.05, **p<0.01, and ***p<0.001 significant in genotype comparison; ‘ns’ = not significant.

**Figure S2.**
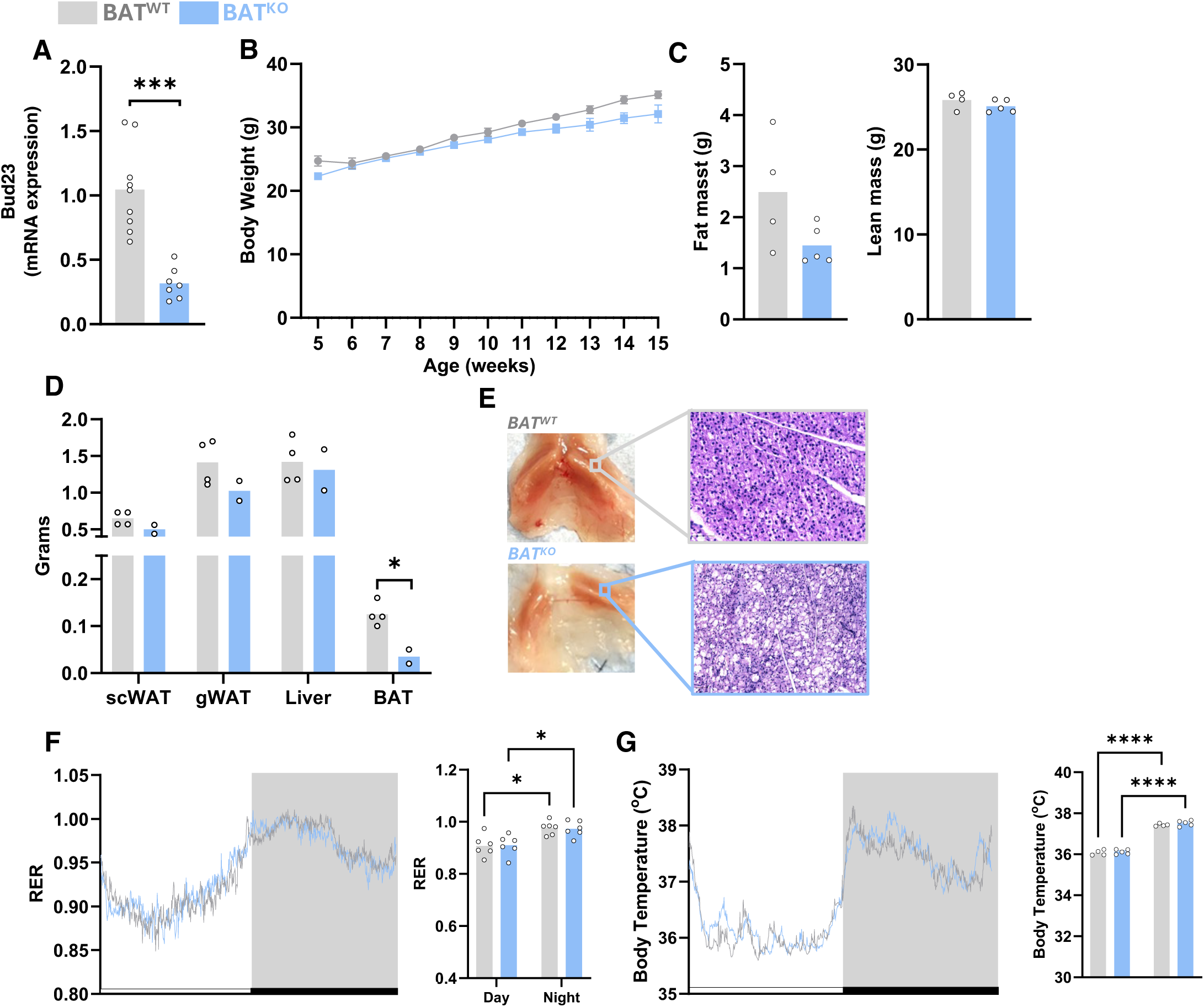
Loss of BUD23 in only in brown adipocytes does not lead to tissue whitening. (**A**) Relative Bud23 expression in BAT of BAT^WT^ and BAT^KO^ mice. (**B, C**) Body weight profile (**B**) and fat and lean mass (**C**; at 15 weeks of age) of BAT^KO^ and BAT^WT^ mice. (**D**) Individual tissue weights of BAT^WT^ and BAT^KO^ mice. (**E**) Representative picture and histology of the BAT from BAT^KO^ and BAT^WT^ mice. (**F, G**) Daily profiles and day/night averages of respiratory exchange ratio (RER; **F**) and body temperature (**G**). *n* reflects biological replicates (mice) in all panels. Mean values are plotted (with ± SEM where appropriate). Statistical significance was tested by unpaired two-tailed Student’s t test (A, C, D) or two-way ANOVA with Holm-Šídák post hoc test (B, F, G). ∗p<0.05, ∗∗∗p<0.001, ∗∗∗∗p<0.0001.

**Figure S3.**
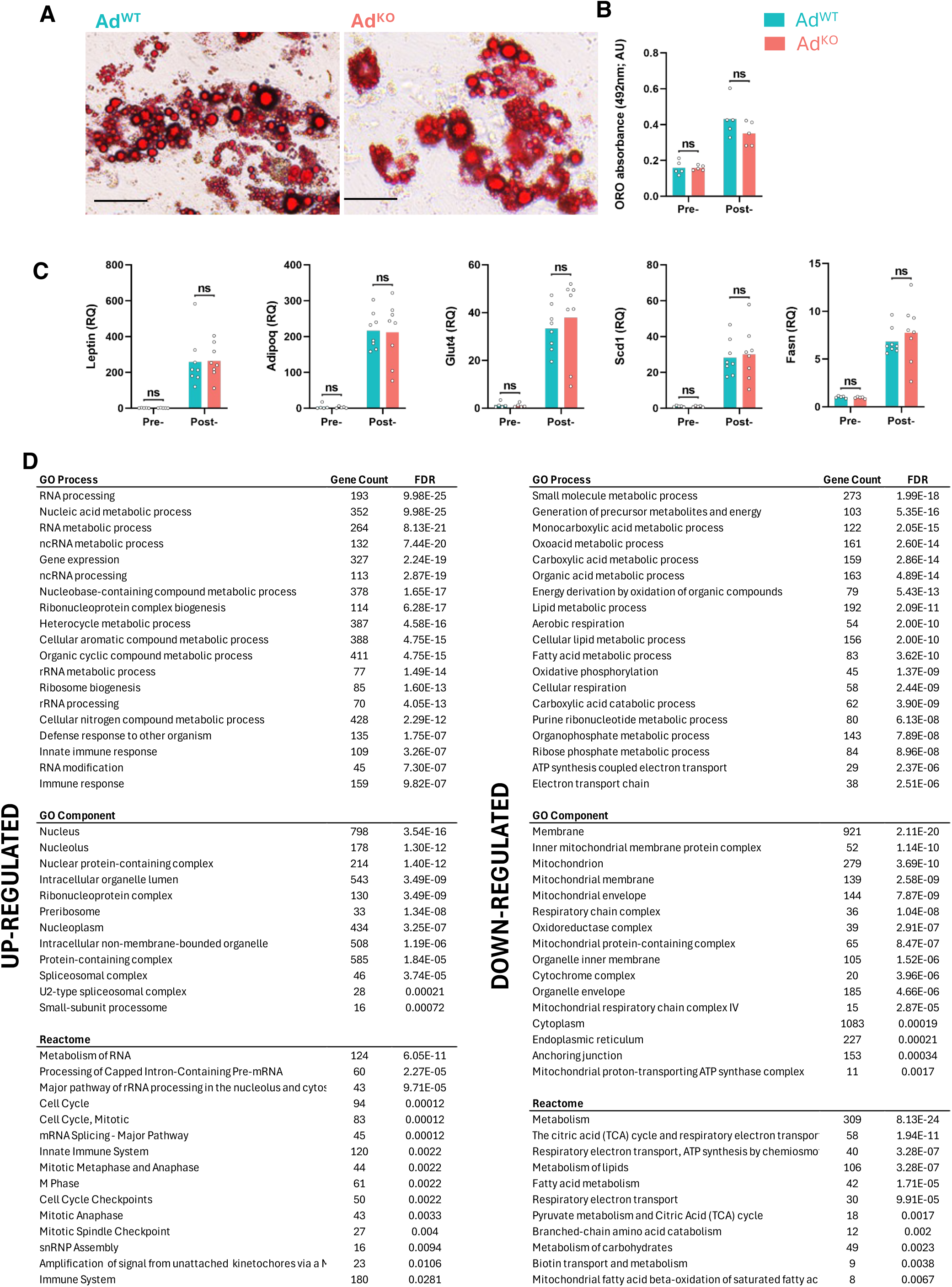
Bud23 is not required for adipocyte differentiation. (**A, B**) Representative Oil Red O (ORO) staining (**A**) and quantification (**B**) in primary adipocytes derived from gWAT of Ad^WT^ and Ad^KO^ mice. (**C**) Relative mRNA expression levels of endocrine hormones, lipolytic and lipogenic genes in gWAT pre-adipocytes from Ad^WT^ and Ad^KO^ mice, pre– and post-differentiation. (**D**) Gene Ontology (GO) terms defined by Biological Process, Cellular Component, and Reactome pathway analysis based on differential gene expression in gWAT of Ad^KO^ and Ad^WT^ mice (13 weeks of age). GO term and Reactome analyses were conducted separately on down-regulated and up-regulated differentially expressed genes (Ad^KO^ vs Ad^WT^). The number of differentially expressed genes associated with each pathway, and the corresponding false discovery rate (FDR) are displayed. Mean values are plotted in **B** and **C**. Statistical significance was tested by two-way ANOVA with Holm-Šídák post hoc test; ‘ns’ = p>0.05.

**Figure S4.**
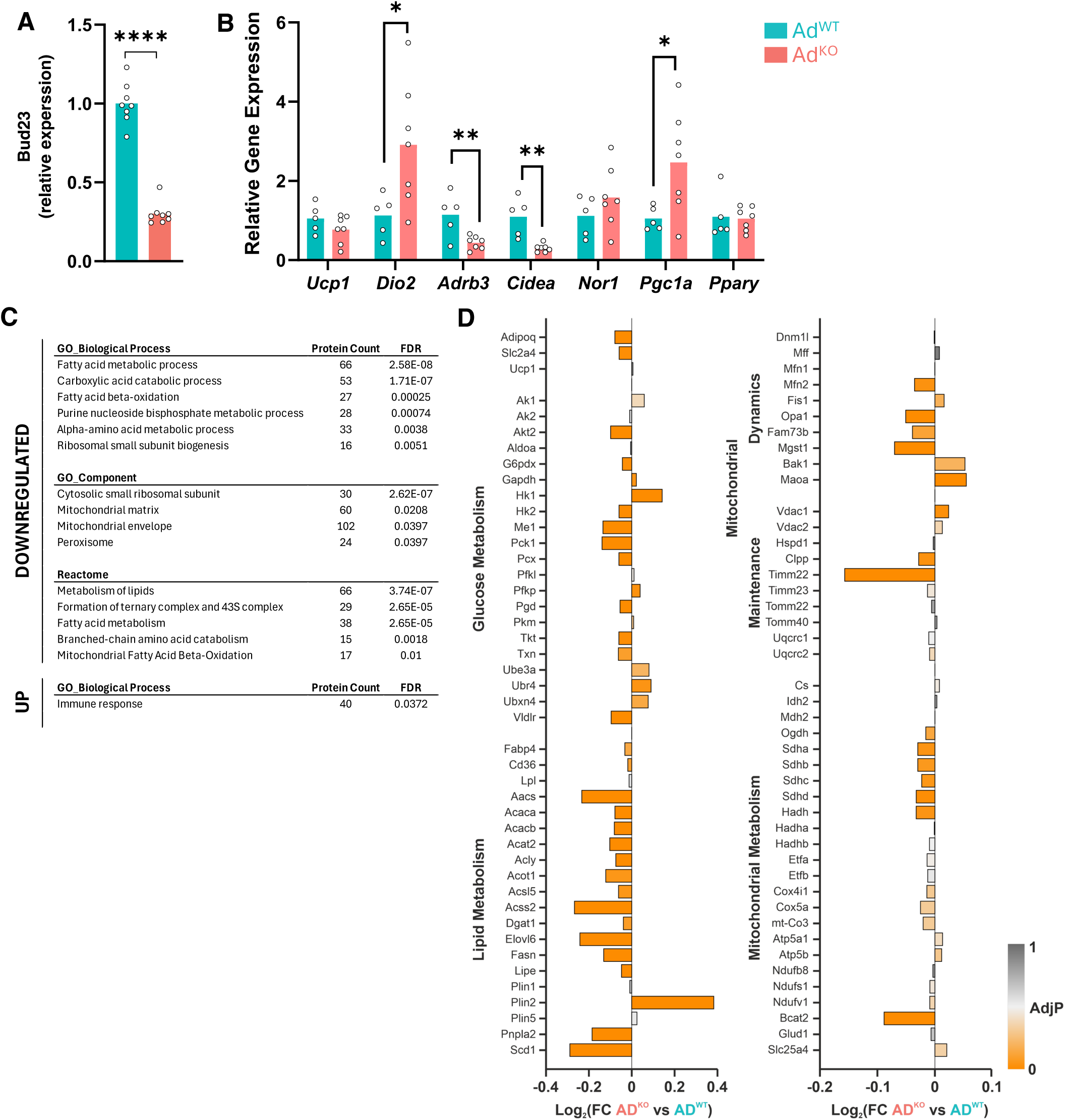
Bud23 directs BAT lipid metabolism and mitochondrial function. (**A**) *Bud23* expression in BAT of Ad^WT^ and Ad^KO^ mice. (**B**) Relative mRNA expression of thermogenic genes in BAT from Ad^WT^ and Ad^KO^ mice (13-weeks of age) housed under standard ambient temperatures (22±1°C). Statistical significance was tested by unpaired two-tailed Student’s t test; ∗p < 0.05, ∗∗p < 0.01, ∗∗∗∗p < 0.0001. Mean values are plotted. (**C**) Gene Ontology (GO) terms defined by Biological Process, Cellular Component, and Reactome pathway analysis based on differential protein expression in Ad^KO^ and Ad^WT^ mice. GO term and Reactome analyses were conducted separately on down-regulated and up-regulated differentially expressed genes (Ad^KO^ vs Ad^WT^). The number of differentially expressed proteins associated with each pathway, and the corresponding false discovery rate (FDR) are displayed. (**D**) Specific protein expression changes (Log_2_FC) from BAT of Ad^KO^ vs Ad^WT^ mice.

**Figure S5.**
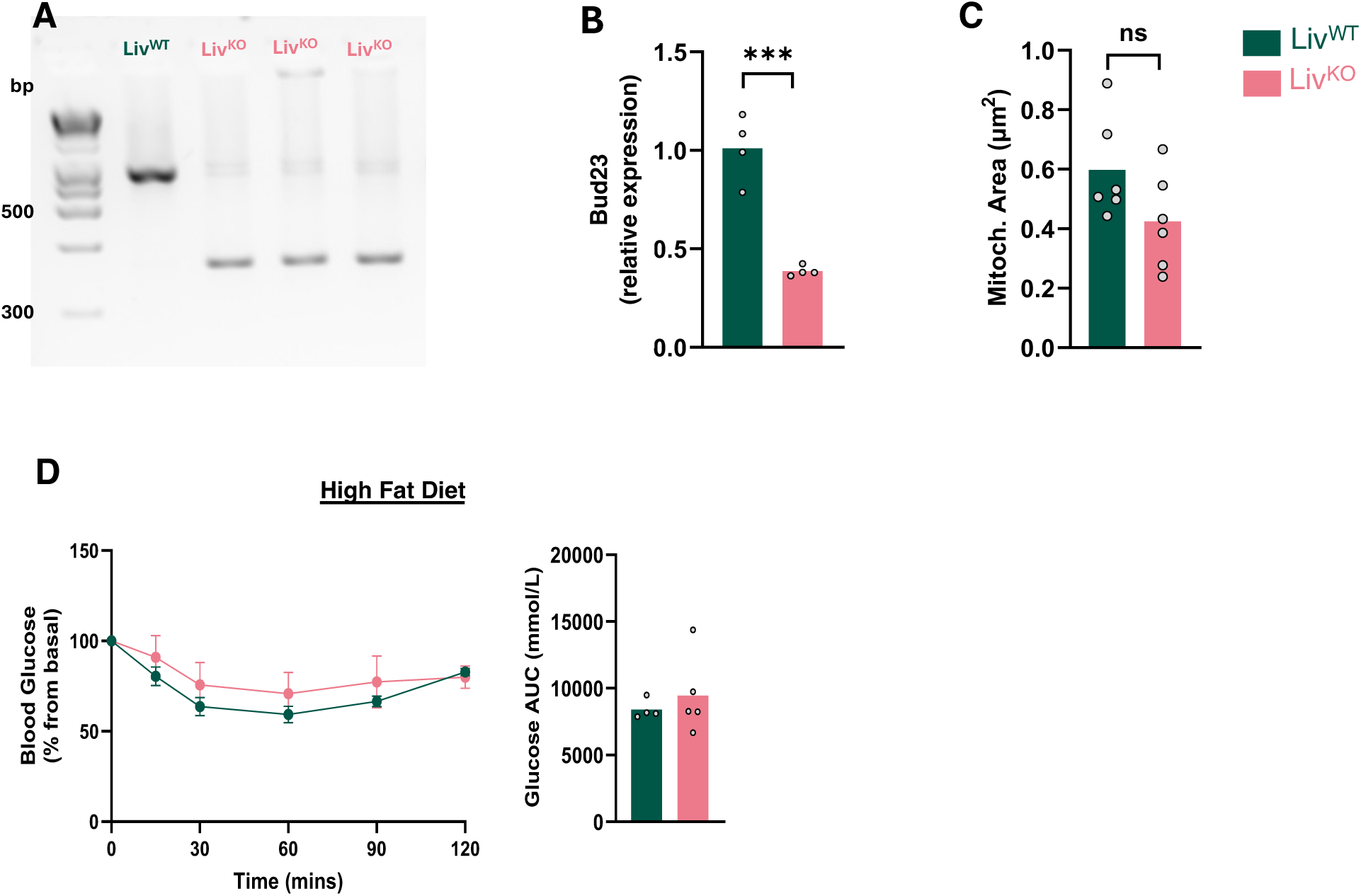
*Bud23* targeting in liver. (**A, B**) Representative agarose gel image of the recombination to ensure knockout (**A**) and relative *Bud23* expression (**B**) in liver of Liv^WT^ and Liv^KO^. (**C**) Liver mitochondrial area assessed on EM images collected from Liv^WT^ and Liv^KO^ mice. (**D**) Glucose levels during insulin tolerance test and area within the curve (AUC) Liv^WT^ and Liv^KO^ mice following 8 weeks of high fat diet (HFD) feeding. Mean values are plotted. Statistical significance was tested by unpaired two-tailed Student’s t test or two-way ANOVA with Holm-Šídák post hoc test; ∗∗∗p<0.001.

**Figure S6.**
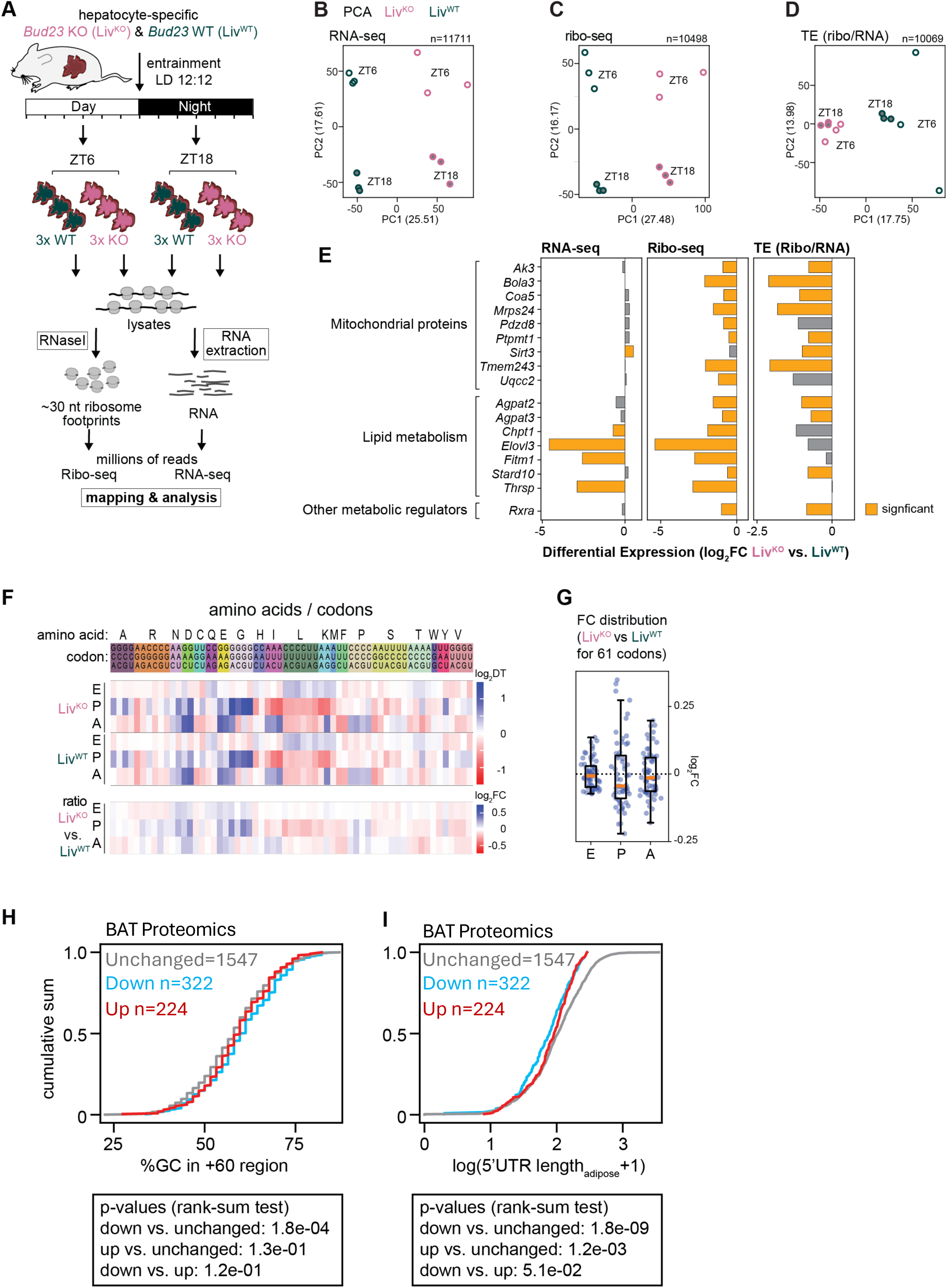
Liver Ribo-seq study design and analyses. (**A**) Schematic showing the design of the ribosome profiling experiment. (**B**) Principal component analysis on the RNA-seq data, showing separation on PC1 by genotype – Liv^KO^ (pink) vs. Liv^WT^ (dark green) – and by timepoint on PC2. (**C**) As in (B) for the ribosome footprint data. (**D**) As in (B) for translation efficiencies (i.e., ratio of normalized Ribo-seq to RNA-seq reads per gene). In this analysis, main separation occurs by genotype (PC1) with little influence by ZT. (**E**) RNA-seq, Ribo-seq and TE log_2_ fold-changes for mitochondrial/metabolism example genes. Significance according to the applied DESeq analysis (see Methods). *Ak3*, involved in nucleotide homeostasis; *Bola3*, involved in mitochondrial iron–sulfur cluster assembly; *Coa5*, cytochrome c oxidase assembly factor; *Mrps24*, mitochondrial ribosomal protein; *Pdzd8*, ER–mitochondria tether, lipid and Ca²⁺ exchange; *Ptpmt1*, mitochondrial inner membrane phosphatase, regulates cardiolipin synthesis; *Sirt3*, mitochondrial NAD⁺-dependent deacetylase; *Tmem243*, mitochondrial transmembrane protein, complex IV assembly; *Uqcc2*, complex III assembly factor. *Agpat2*, 1-acylglycerol-3-phosphate O-acyltransferase, phospholipid synthesis; *Agpat3*, same enzyme family as above, phospholipid metabolism; *Chpt1*, choline phosphotransferase, phosphatidylcholine synthesis; *Elovl3*, elongation of very long chain fatty acids; *Fitm1*, fat storage–inducing transmembrane protein, lipid droplet formation; *Stard10*, lipid transfer protein (phospholipids, phosphoinositides); *Thrsp*, thyroid hormone–responsive protein (Spot14), regulator of lipogenesis. *Rxra*, retinoid X receptor alpha, transcription factor for lipid/energy metabolism. (**F**) Analysis of dwell times of elongating ribosomes using the RiboDT pipeline {Gobet, 2022 #2092}. Upper two sets show heat-maps of dwell times for E-P– and A-sites in Liv^KO^ and Liv^WT^ data (ZT6 and ZT18 combined data) and indicate very similar preference for codon occupancies between genotypes. Lower part shows fold-change differences in dwell times, Liv^KO^ vs. Liv^WT^, again indicating relatively mild differences in codon-specific dwelling between genotypes. (**G**) Box plots of dwell time log_2_ fold changes between genotypes (same as lower panel of (F)) at E-, P– and A-sites indicate that overall codon-specific variation appears more prominent at the P-site as compared to E-or A-sites. (**H**) Analysis of association of the features identified for liver in Figure 6 with protein expression changes identified by proteomics in BAT. In blue, the 322 proteins with significantly lower abundance in BAT display increase GC content in the 60 nt downstream of the translational start site. Red indicates significantly upregulated proteins (N = 224) and grey the proteins with unchanged abundance (N = 1547). (**I**) As in (H), but for the feature “short 5’ UTR” that was also identified from the liver Ribo-seq data in Figure 6. Comparison with BAT proteomics data shows association of downregulated proteins with shorter UTRs. For the analysis, adipocyte-specific CAGE data was used (ST2 mesenchymal stem cells, differentiation to adipocytes, RIKEN).

**Figure S7.**
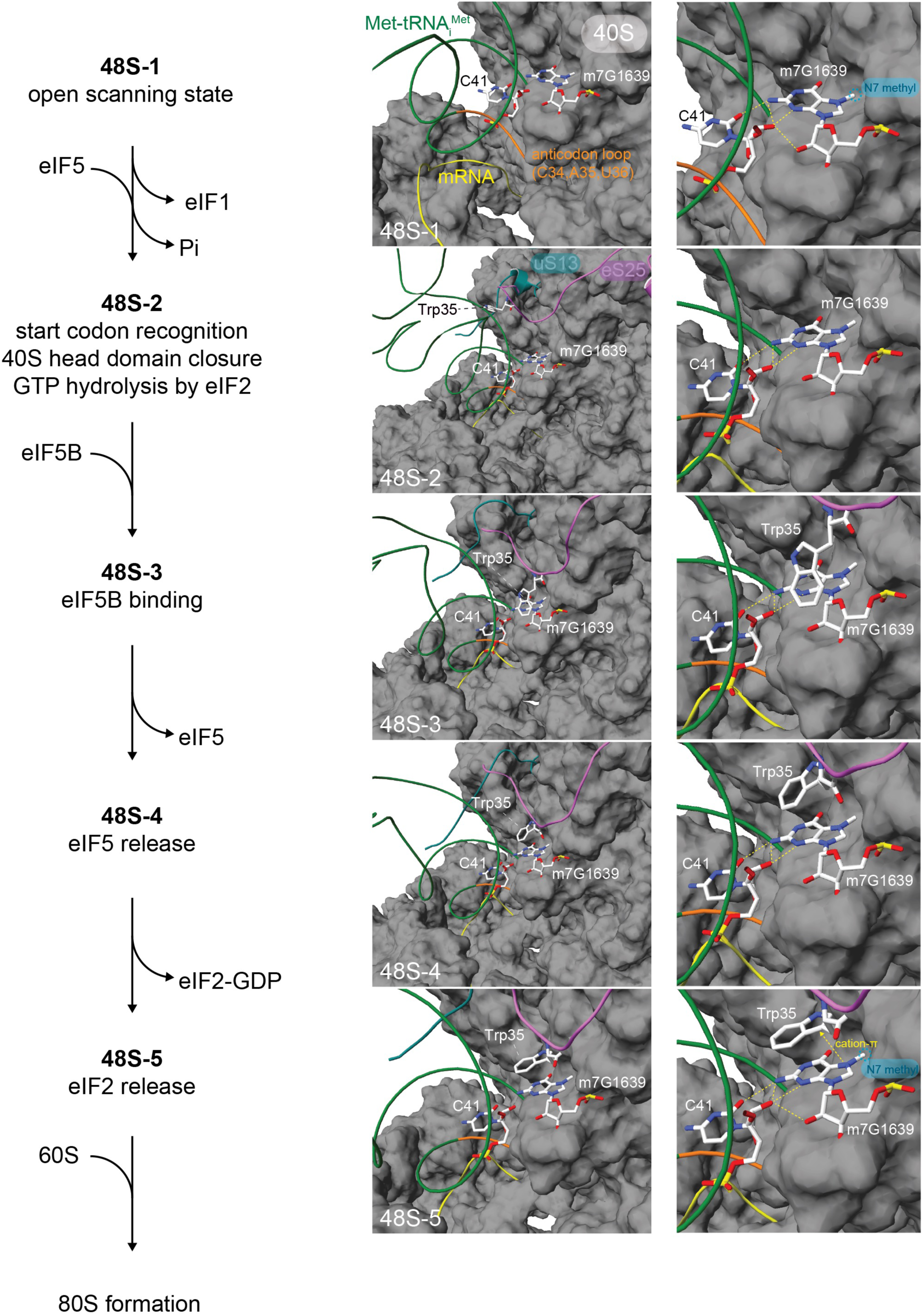
Contacts of m^7^G1639 in initiating ribosome structure model. Overview of structural models of human early (48S-1) to late (48S-5) initiation complexes emphasizing m^7^G1639 surroundings (based on PDB structures 8pj1-5). Left part of the figure illustrates the sequence of 48S state transitions to 80S formation with major rearrangements between them. Middle panels are zoom-out, and right panels zoom-in views of the structural models. Important highlighted residues apart from m^7^G1639 (with N^7^-methyl group marked by blue halo in some of the panels) are Met-tRNA_i_^Met^ (backbone traced in dark green; anticodon loop in orange), tRNA residue C41 that forms extensive hydrogen bonds with m^7^G1639; eS25/RPS25 (backbone traced in purple) Trp35 is progressively brought close to m^7^G1639 up to a point where it makes cation-π stacking onto m^7^G1639 in the last structure). The C-terminal region of 40S ribosomal protein RPS18/uS13 backbone is visible as a teal ribbon, mRNA trace is marked in yellow and 18S rRNA surface (excluding m7G1639 to see its interactions) in grey.

**Figure S8.**
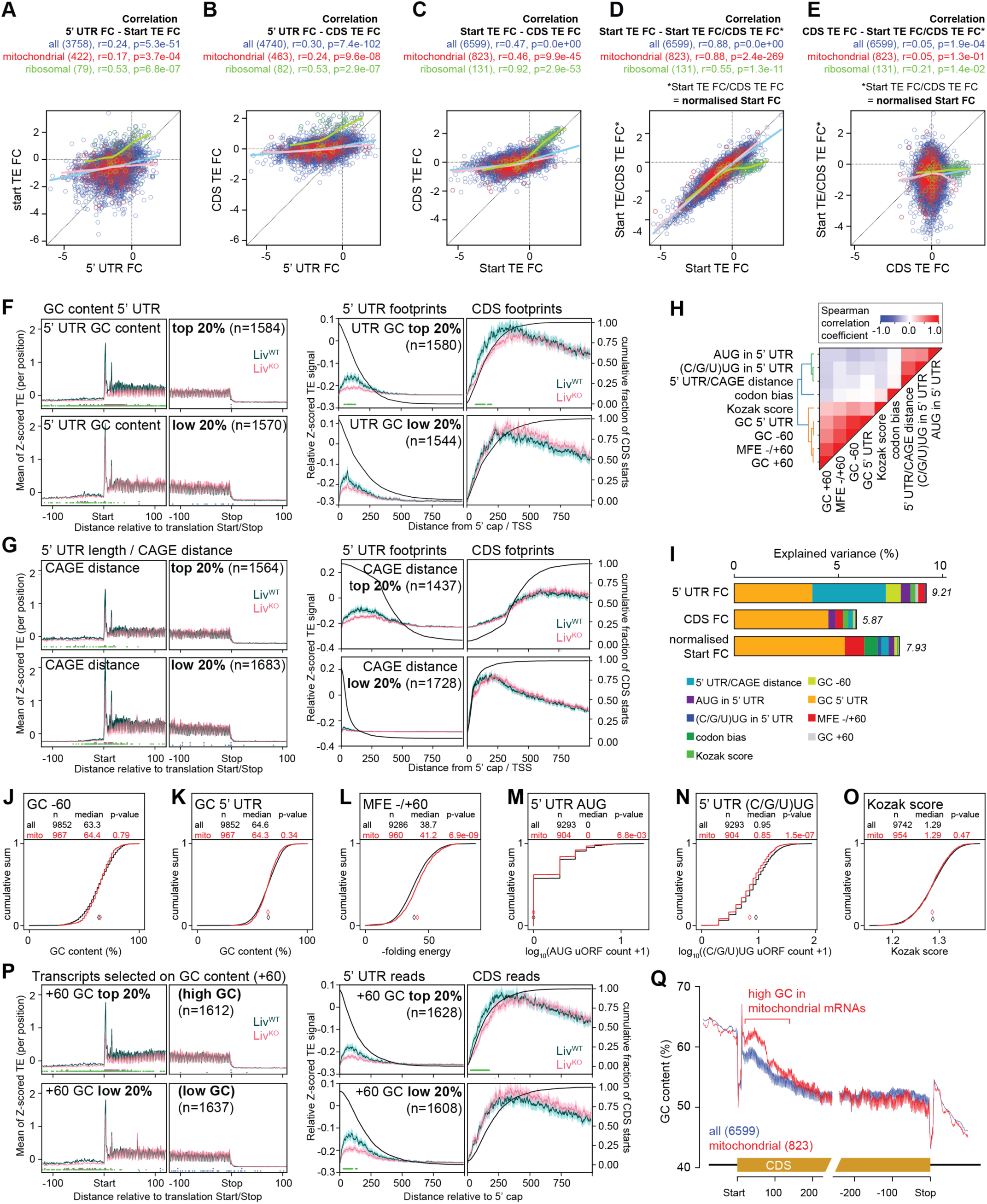
Further analysis of predictors of BUD23-sensitivity. (**A**) Complementary to Fig. 6B, the graph shows a correlation analysis between 5’ UTR TE fold-change and Start TE fold-change, for all (blue), mitochondrial (red) and ribosomal (green) transcripts. The local regression fit (LOESS) is shown as line for each transcript group. Spearman correlation coefficient and p-value of correlation are given above, alongside number of transcripts used in the analysis. (**B**) As in (A) for correlation of 5’ UTR TE fold-change and CDS TE fold-change. (**C**) As in (A) for correlation of Start TE fold-change and whole CDS TE fold-change. (**D**) As in (A) for correlation of Start TE fold-change and Start TE fold-change relative to whole CDS TE fold-change. (**E**) As in (A) for correlation of Start TE fold-change (corrected for CDS TE fold-change) and CDS TE fold-change. (**F**) Left panels: Metagene plot aligning ribosome footprint A-sites relative to coding sequence (CDS) around start and stop codons, for Liv^KO^ (pink) and Liv^WT^ (dark green) for transcripts with specifically high (upper) and low (lower) GC content in 5’ UTR. Right panels: Alignment of reads to the annotated transcriptional start site (TSS), and plotted separately for reads that fall into annotated 5’ UTR (left) and CDS (right). Black line shows cumulative density of 5’ UTRs (left) and CDS starts (right). (**G**) As in (F), but for transcripts with particularly long (upper) or short (lower) 5’ UTRs according to the CAGE data. (**H**) Cross-correlation analysis between transcript sequence features considered in the analysis. (**I**) Linear regression model’s prediction of TE FC variances explained by different transcript sequence features (color-coded; listed at bottom). TE FCs are predicted for 5’ UTRs, CDSs and Start regions (first 20 codons, relative to whole CDS FCs). Total explained variance by a combination of transcript sequence features is given at the right of the bars. (**J**) Analysis of GC content within the 60 nt upstream of the initiation codon, for transcripts encoding mitochondrial proteins (red) vs. all transcripts (black), indicating no significant differences in GC-content in the mitochondrial group. In the upper part of the panel, the number of transcripts used for the analysis (n), their median (also shown as diamond symbol in graph) and the p-value for the difference from a ranksum test are given. (**K**) As in (J), for GC content of the full 5’ UTR, for transcripts encoding mitochondrial proteins (red) vs. all transcripts (black). (**L**) As in (J), for Minimal Free Energy (MFE) of RNA folding in the window +/-60 nt around the CDS initiation codon. (**M**) As in (J) for AUG count within the 5’ UTR (and taking into account ribosome footprint coverage, serving as a proxy for AUG-initiated uORFs). (**N**) As in (M), but for alternative uORF start codons, (C/G/U)UG. (**O**) As in (J) for Kozak score. (**P**) As in (F), but for transcripts with particularly high (upper) or low (lower) GC content in the +60 nt region. (**Q**) Percentages of all (blue) and of mitochondrial protein transcripts (red) with G or C nucleotides at positions around the start and stop codons. A 10 nt moving window was used for averaging to smoothen the curves. It is evident that mitochondrial transcripts show higher GC content at the CDS beginning as compared to all transcripts.

**Figure S9.**
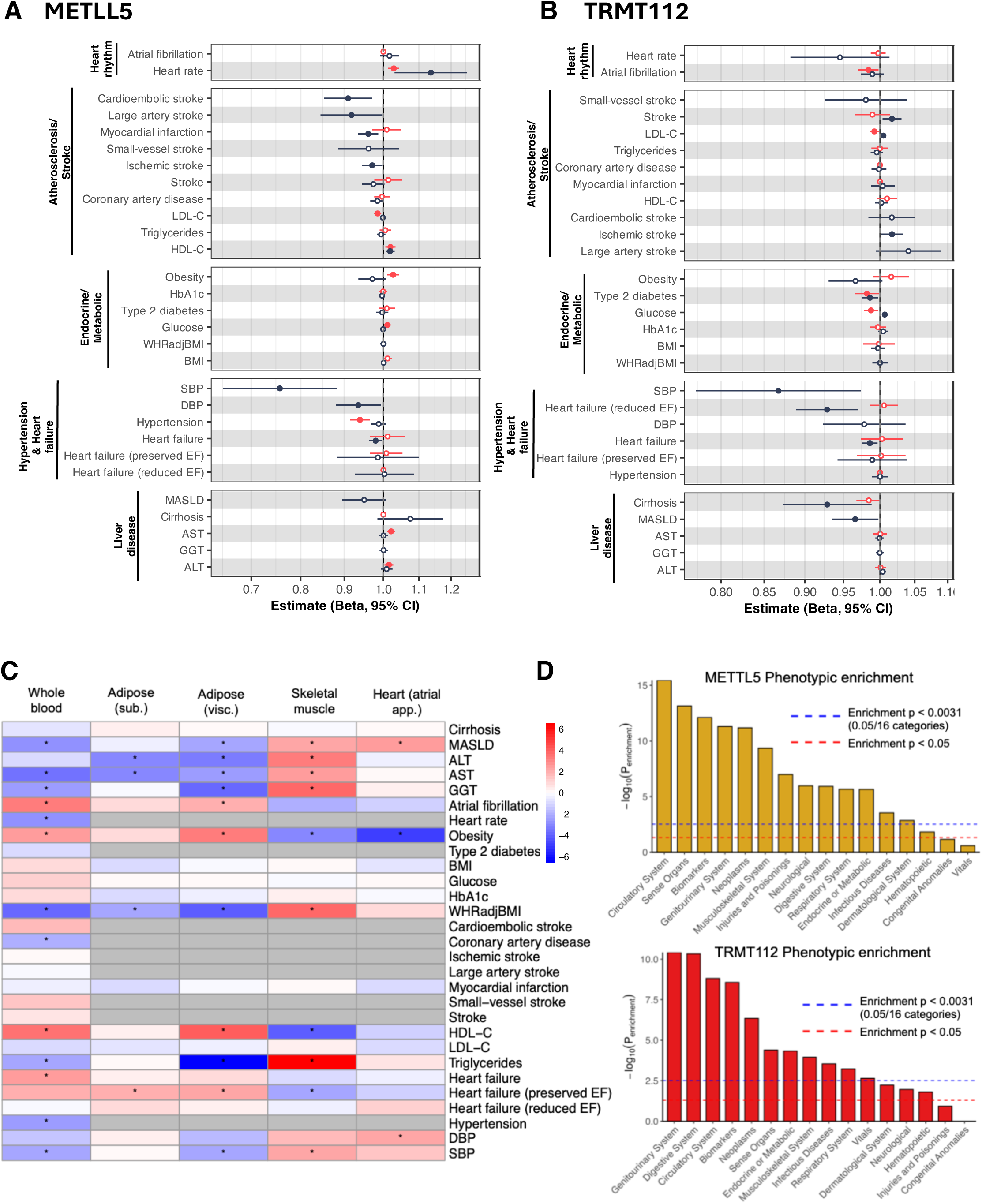
Mendelian randomization analysis of whole blood METTL5 and TRMT112 expression across cardiometabolic traits and tissues. This figure presents Mendelian Randomization (MR) analyses examining the association of genetically predicted expression of METTL5 and TRMT112 in whole blood with cardiometabolic outcomes. (**A**) shows forest plots of MR estimates (Beta ± 95% CI) for METTL5 expression across primary and replication GWAS datasets. (**B**) presents the corresponding MR results for TRMT112. (**C**) displays a heatmap comparing the Z-scores (Beta/SE) for BUD23 across cardiometabolic tissues, including whole blood, subcutaneous and visceral adipose tissue, skeletal muscle, and heart atrial appendage. Asterisks (*) denote nominal significance (P < 0.05), and grey cells indicate unavailable instruments. (**D**) shows enrichment analyses for METTL5 and TRMT112, respectively, across predefined clinical categories in the Million Veteran Program (MVP). Bar heights reflect enrichment significance, and domains meeting Bonferroni or nominal thresholds are highlighted. All effect directions correspond to associations per increased gene expression in whole blood.

